# Improving protein expression, stability, and function with ProteinMPNN

**DOI:** 10.1101/2023.10.03.560713

**Authors:** Kiera H. Sumida, Reyes Núñez-Franco, Indrek Kalvet, Samuel J. Pellock, Basile I. M. Wicky, Lukas F. Milles, Justas Dauparas, Jue Wang, Yakov Kipnis, Noel Jameson, Alex Kang, Joshmyn De La Cruz, Banumathi Sankaran, Asim K. Bera, Gonzalo Jiménez-Osés, David Baker

## Abstract

Natural proteins are highly optimized for function, but are often difficult to produce at a scale suitable for biotechnological applications due to poor expression in heterologous systems, limited solubility, and sensitivity to temperature. Thus, a general method that improves the physical properties of native proteins while maintaining function could have wide utility for protein-based technologies. Here we show that the deep neural network ProteinMPNN together with evolutionary and structural information provides a route to increasing protein expression, stability, and function. For both myoglobin and tobacco etch virus (TEV) protease, we generated designs with improved expression, elevated melting temperatures, and improved function. For TEV protease, we identified multiple designs with improved catalytic activity as compared to the parent sequence and previously reported TEV variants. Our approach should be broadly useful for improving the expression, stability, and function of biotechnologically important proteins.

## Introduction

Evolution has optimized function over stability in many natural proteins^1^; as a result, they often exhibit poor solubility, thermostability, and expression in heterologous systems, all of which reduce the yield of functional protein.^2,3^ Many protein-based therapeutics and catalysts are limited in their industrial application by low stability, making protein stabilization a research area of increasing interest.^4,5^ Experimental methods such as directed evolution have been extensively used to optimize desirable features in proteins, but are often prohibitively resource- and labor-intensive.^6,7^ Computational tools have been developed to achieve the benefits of directed evolution while minimizing experimental screening.^8–11^ PROSS (protein repair one-stop shop), for example, utilizes evolutionary information and Rosetta physics-based energy calculations to perform sequence redesign using a three-dimensional (3D) structure as input, and has been shown to increase soluble expression and thermostability of several natural proteins.^8^ More recently, advances in deep learning-based modeling of proteins have been applied to generate new variants of natural proteins, including language models that generate sequences for a given enzyme family or function^11^, convolutional neural networks that leverage structural information for prediction of gain-of-function mutations^10^, and shallow neural networks for guiding combinatorial directed evolution^12^.

Deep learning-based tools for protein sequence design have shown success in the generation of novel proteins with excellent expression, solubility, and sub-angstrom accuracy to design models.^11,13,14^ ProteinMPNN generates highly stable sequences for designed backbones, and for native backbones, generates sequences that are predicted to fold to the intended structures more confidently than their native sequences.^13^ We reasoned that ProteinMPNN could be applied to protein stability optimization, and set out to develop a strategy for applying ProteinMPNN to natural proteins to increase solubility and stability. We chose as model systems one of the first proteins whose structure was solved, the oxygen storage protein myoglobin, and the widely used protease from tobacco etch virus (TEV).

### Protein stabilization with ProteinMPNN

ProteinMPNN generates amino acid sequences that are predicted to fold to a given 3D structure. The method is purely structure-based, and does not have access to functional information. Therefore, to retain protein function during sequence design, additional information must be provided to the network. We experimented with a range of approaches to retain functionality during the design process. In all targets, to preserve the catalytic machinery and substrate-binding site, we fixed the amino acid identities of the first shell functional positions – defined as those within 7 Å of the substrate in a ligand-bound crystal structure complex. In TEV protease, we used evolutionary information to further identify residues critical to activity. In myoglobin, we performed limited backbone redesign to further stabilize the structure. With the design space selected, we performed sequence design with ProteinMPNN, predicted the structures with AlphaFold2^15^, and filtered by predicted local distance difference test score (pLDDT) and Cα root mean square deviation (RMSD) to the input structure (Figure 1).

**Figure 1.**
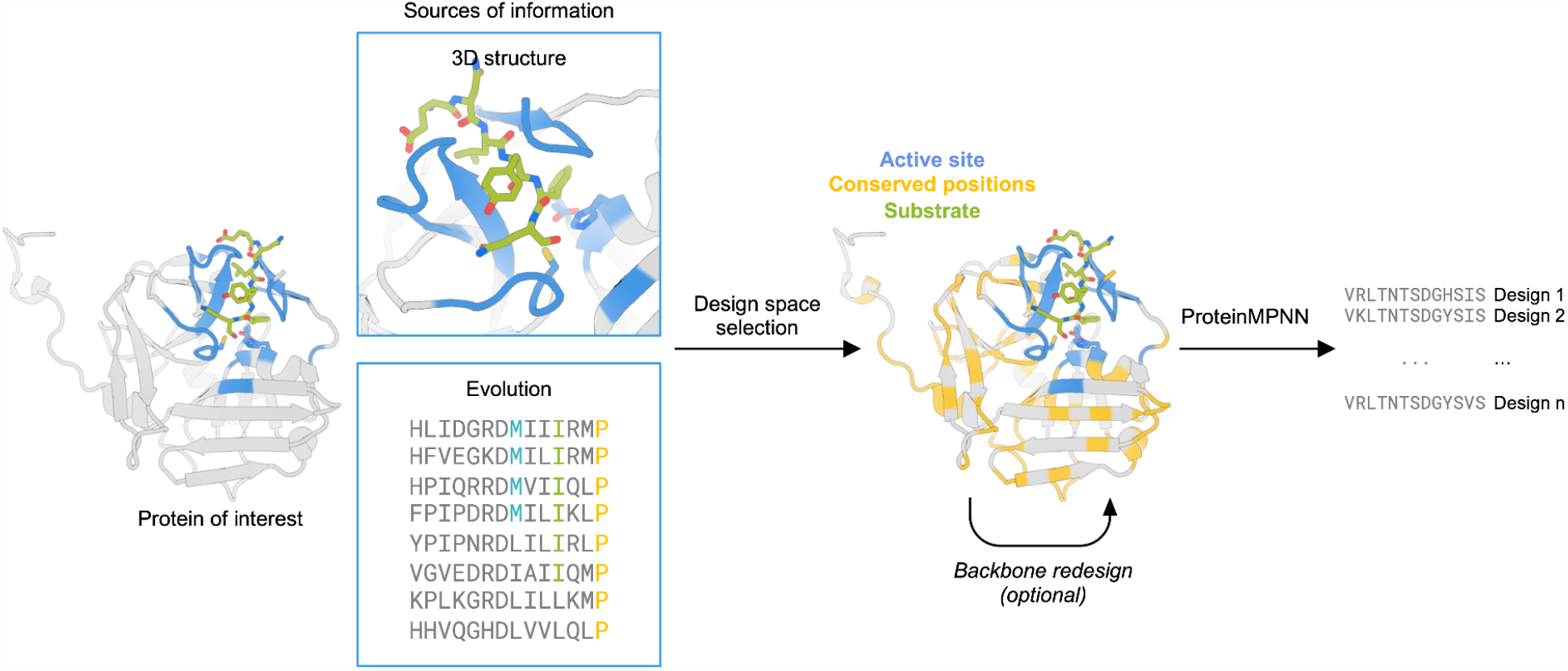
Design strategy for optimization of protein expression and stability using ProteinMPNN. The design space is chosen to preserve native protein function by fixing the amino acid identities of residues close to the ligand/substrate and those that are highly conserved in multiple sequence alignments. The protein backbone structure and fixed position information of amino acids are input into ProteinMPNN, which generates new amino acid sequences likely to fold to the input structure. The backbone structure in loop regions can optionally be remodeled using RoseTTAfold joint inpainting to further idealize the input protein.

### Design of myoglobin variants with increased stability

We first applied our design strategy to the model protein myoglobin. Myoglobin binds heme to carry oxygen in mammalian muscle tissue^16^ and has relevance in clinical applications as a biomarker^17^, as a versatile platform for biocatalytic applications^18–20^, and in food science as an ingredient in artificial meat products^21–23^. Current efforts to create more stable variants of myoglobin have focused on the stabilization of the globin fold through stapling with cysteine-reactive noncanonical amino acids.^24,25^

We applied our ProteinMPNN design protocol described above starting from a crystal structure of human myoglobin, nMb (PDB: 3RGK)^26^. To preserve the oxygen storage function, we fixed the identities of 17 positions located around the heme ligand in the heme-bound structure (Figure 2A). 60 sequences were generated with ProteinMPNN and evaluated for their likelihood to recapitulate the myoglobin backbone coordinates using AlphaFold2 single-sequence predictions (see methods). Eight of the designs did so with high confidence (pLDDT > 85.0 and Cα RMSD < 1.0; analogous single-sequence prediction of the native sequence yielded pLDDT = 50.6 and Cα RMSD = 7.5). Four designs with close structural agreement in the heme-binding region were selected for experimental testing.

**Figure 2.**
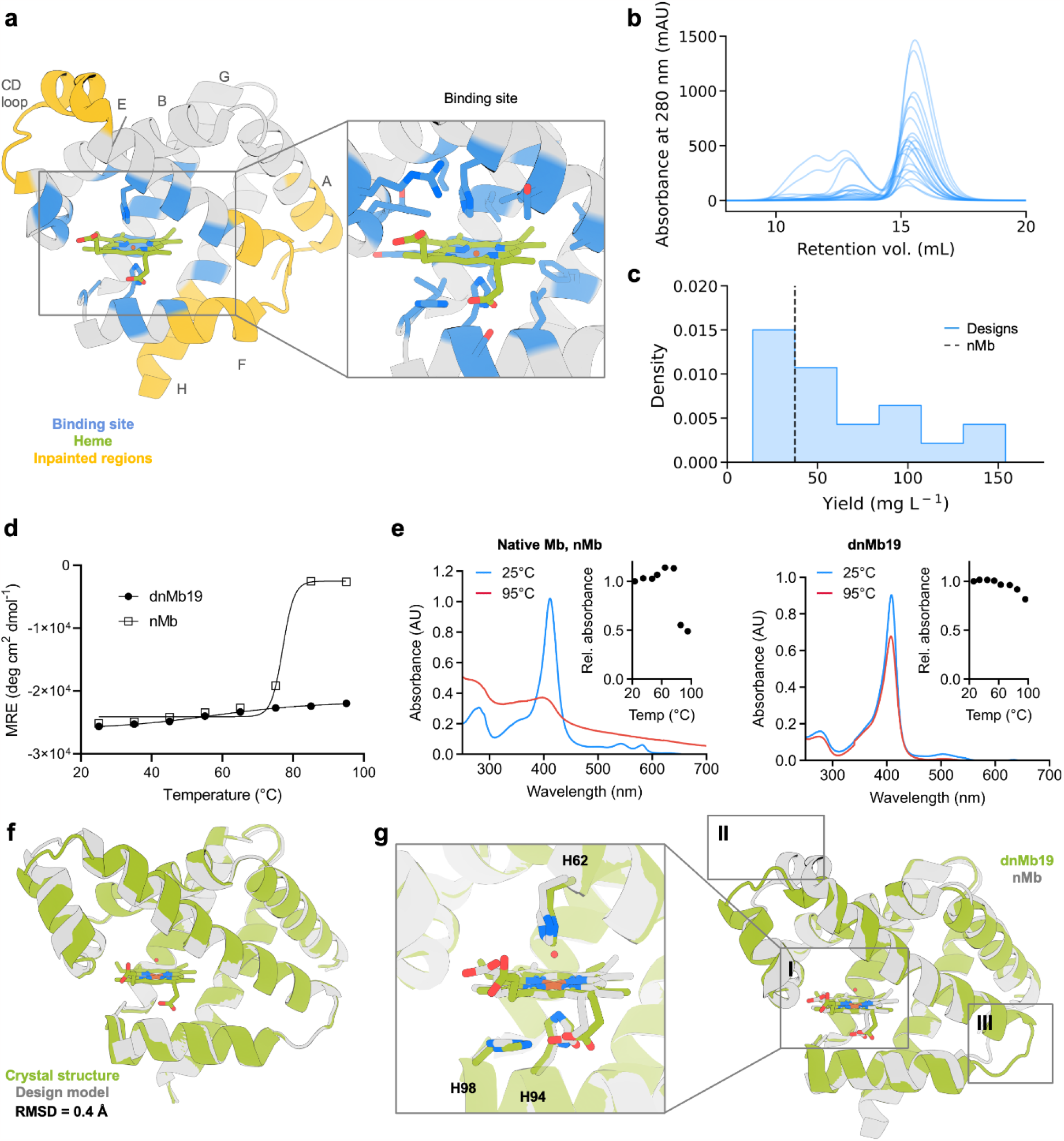
ProteinMPNN design improves myoglobin expression and thermostability. (A) Positions adjacent to the heme were kept fixed during sequence design (shown in blue). Non-conserved regions (in yellow) were subjected to backbone remodeling. Inset shows the heme-binding site. (B) SEC traces of 144 designed myoglobin variants. (C) Soluble yield of myoglobin designs and native myoglobin nMb (represented as a black dashed line). (D) CD melting temperature plots of dnMb19 compared to native myoglobin (signal reported in molar residue ellipticity (MRE)). (E) Absorbance plots of dnMb19 and native myoglobin (inset shows temperature scan). (F) Structural alignment of the crystal structure (green) and AlphaFold2 (AF2) prediction (gray) of dnMb19. (G) Overlay of the crystal structure of native myoglobin (gray) and the crystal structure of dnMb19 (green, PDB: 8U5A). Non-conserved regions displayed in insets II and III were subjected to backbone redesign.

We also explored limited backbone redesign of poorly ordered regions to attempt to further stabilize the protein. The globin superfamily, of which myoglobin is a member, has a fold made up of eight alpha helical regions, with diversity in the termini and two loop regions flanking the heme-binding pocket^27–29^ (Figure S1). We selected these less-conserved loop regions for backbone remodeling with RoseTTAFold joint inpainting (Figure 2A).^30^ We generated two distinct sets of designs with structural remodeling: one with the region joining helices E and F redesigned, and one additionally including the CD-loop region (Figure 2A). From these remodeled backbones, we again performed sequence design with ProteinMPNN with the heme-binding site kept fixed as described above. Following filtering on structure prediction metrics (Figure S2), 20 sequences were selected for experimental testing. All 20 designs have 41-55% sequence identity to the most similar protein (a myoglobin in all cases) in the UniRef100 database^31^ (Table S1).

Synthetic genes encoding the designs and the parent sequence, nMb, were expressed in *E. coli*. The heme-loaded *holo*-proteins were purified via immobilized metal affinity chromatography (IMAC) and size exclusion chromatography (SEC). All designs were solubly expressed and monomeric by SEC (Figure 2B). 13 of the 20 designs had higher levels (up to 4.1-fold increase) of total soluble protein yield (Figure 2C). All 20 designs had similar heme-binding spectra to native myoglobin, with agreement in Soret maximum (407-413 nm vs 409 nm in native) and Q band features (500, 537, 582 and 630 nm), suggesting preservation of the native heme-binding mechanism (Figure S3).

The thermal stabilities of eight highly-expressing designs (six and two designed with and without backbone remodeling, respectively) were evaluated using circular dichroism (CD) spectroscopy. All eight designs had higher melting temperatures than native myoglobin, with six remaining fully folded at 95 °C (native myoglobin melts at 80 °C; Figures 2D and S4). Heme binding was also evaluated over a temperature gradient to determine functional thermal stability. All designs preserved heme binding at higher temperatures than native myoglobin (as monitored by changes to Soret band wavelength and intensity in the UV/Vis spectrum), with five designs maintaining significant heme-binding at 95 °C (Figure S5). One of the five designs, dnMb19, generated with the more aggressive backbone remodeling strategy, showed much higher thermal stability of heme binding compared to native myoglobin (Figure 2E). Overall, remodeling regions of the myoglobin backbone with inpainting increased the success rate for retaining heme-binding at elevated temperatures.

To understand the structural basis of these improvements in stability, we solved the crystal structure of dnMb19 (2.0 Å resolution, PDB: 8U5A). We found that it closely agreed with the AlphaFold2 prediction (0.4 Å Cα RMSD, Figure 2F), including the regions remodeled with inpainting. Native sidechain contacts with the heme group are largely preserved in dnMb19 (Figure 2G, inset I). Outside of the heme-binding site, the crystal structure confirms the structural changes introduced by inpainting: the C and E helices were elongated as designed and connected by a new loop (Figure 2G, inset II); the loop connecting the E and F helices has a new conformation, and the F helix was straightened through the replacement of PRO88 with GLU89 (Figure 2G, inset III). These results illustrate the power of RoseTTAFold joint inpainting and ProteinMPNN to accurately remodel native protein backbones while increasing solubility, thermostability, and functional stability.

### Design of TEV protease variants with improved stability and catalytic activity

To explore the utility of ProteinMPNN sequence design for stabilizing enzymes, we next applied our design strategy to the cysteine protease from tobacco etch virus (TEV). TEV protease is widely used in biotechnological applications to specifically cleave between glutamine and serine in its recognition sequence (ENLYFQ/S) to remove purification tags from recombinant proteins. However, it is often difficult to use TEV protease due to its minimal soluble yield, low thermostability, and poor catalytic activity. These properties often necessitate long incubation times and result in incomplete cleavage.^32^

We applied our sequence design strategy to TEV protease starting from an autolysis-resistant S219D variant, TEVd (PDB: 1LVM)^33^. We defined the active site residues as described above to fix during redesign. We additionally fixed the amino acid identities of residues that are most conserved within the protein family (determined from a sequence alignment generated against UniRef30^31^), as residues distant from the active site can contribute significantly to function.^34^ We ranked each amino acid identity at each position by degree of conservation in the sequence alignment and varied the percentage of these most highly conserved residues to fix during sequence redesign between 30-70%. We generated four distinct sets of designs that fixed the amino acid identities of just the active site residues, or the active site residues and 30%, 50%, and 70% of the most conserved residues in the TEV family (Figure 3A, see Methods). 144 sequences were generated with ProteinMPNN which were all predicted with high confidence to fold to the TEV structure by AlphaFold2 (pLDDT > 87.5; native TEV is predicted with pLDDT = 90) and possess 55% to 85% sequence identity to the parent sequence. All 144 designs were selected for experimental testing.

**Figure 3.**
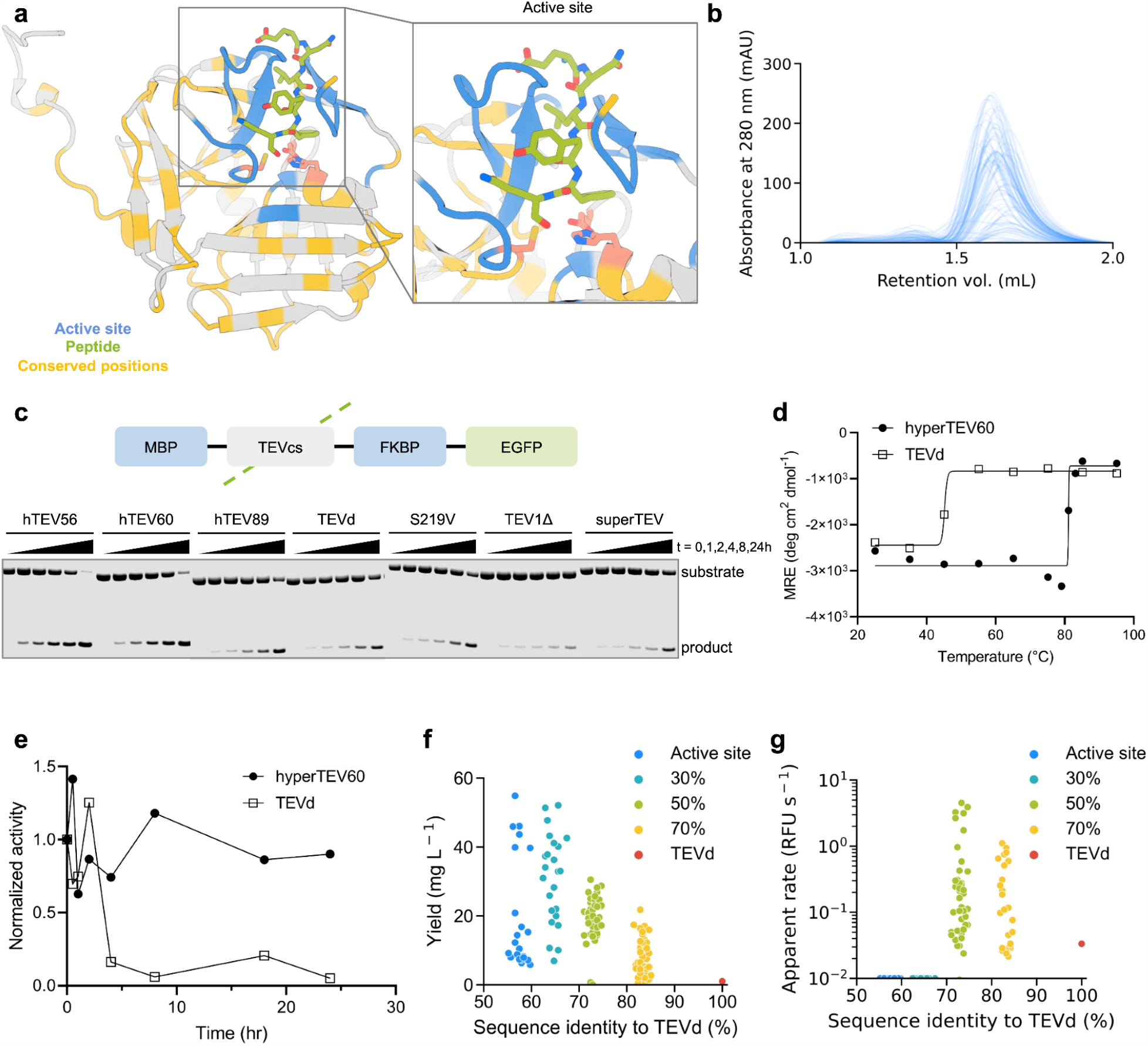
ProteinMPNN sequence design improves TEV protease expression, thermostability, and catalytic efficiency. (A) TEVd (PDB: 1LVM) input structure with positions fixed during redesign highlighted. Active site residues surrounding the substrate (blue), and highly conserved residues (yellow), and catalytic residues (pink) are highlighted. Inset shows zoom-in of the active site region. (B) SEC traces of designed TEV variants. (C) Diagram of TEV substrate (top) and fluorescent gel image of TEV cleavage reactions at various time points (bottom). (D) CD melting temperature plots of designed and native TEV (signal reported in molar residue ellipticity (MRE)). (E) Benchtop stability comparison of native TEVd and designed variant assessed as activity measured over time incubated at 30 °C before inclusion in assay. (F) Decreased evolutionary constraints correlate with higher soluble expression levels. Legend indicates regions fixed during design (all designs have active site fixed). (G) Designs made with the active site and 50% most conserved residues fixed during design exhibited highest catalytic activity. Raw apparent rate reported in relative fluorescence units (RFU) per second.

Synthetic genes encoding the designs, the parent sequence, TEVd, and several previously reported TEV variants were expressed in *E. coli*, and the resultant proteins were purified via IMAC and SEC. 134 of 144 designs expressed solubly and were monomeric by SEC (Figure 3B). 129 of 144 designs exhibited higher levels of soluble expression than TEVd (TEVd average yield = 1 mg/L culture, designs average yield = 20.1 mg / L culture (Figure 3F)).

We evaluated catalytic activity using a previously described^7^ coumarin derivative with 7-amino-4-trifluoromethylcoumarin conjugated to the C-terminus of the substrate peptide Ac-ENLYFQ (Figure S7A). Purified protein was incubated with the peptide-coumarin substrate and 64 designs displayed progress curves with fluorescence above background, indicating substrate turnover (Figures S7B and S7C). Designs made with no evolutionary constraints had improved soluble expression over the parent but were not active on the peptide substrate, while designs with the highest activities were designed with the top 50% most conserved residues fixed (Figures 3F and 3G). We performed detailed kinetic analysis of three highly active designs from the 50% method – hyperTEV56, hyperTEV60, and hyperTEV89 – and the parent sequence TEVd^8^. The designs displayed improved catalytic efficiencies (*k*_cat_/*K*_m_) compared to TEVd, with up to 26-fold improvements (Table 1 and Figure S8).

**Table 1.**
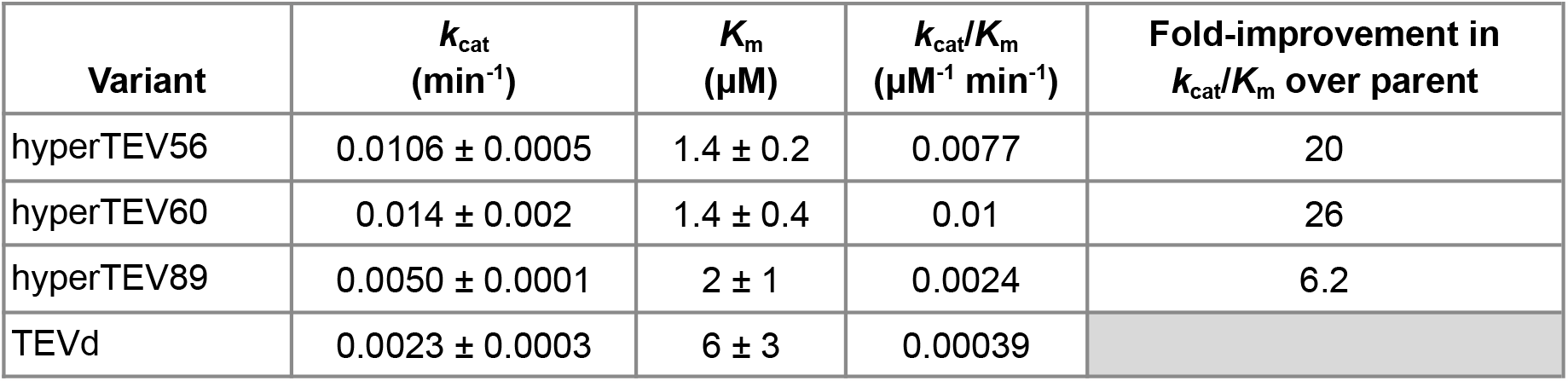
Kinetic parameters for TEV redesigns and parent TEV variant. Kinetic parameters derived from the cleavage assay with the fluorescent peptide-coumarin substrate in Figure S7A. Uncertainties are standard deviations of values calculated from fitting three technical replicates.

Next, we tested the most active designs with a fusion protein substrate to assess performance on the target application of tag removal. The designs and a set of previously engineered TEV proteases^32,33,35–37^ were incubated at 30 °C with the fusion protein substrate MBP-TEVcs-FKBP-EGFP, where MBP is maltose-binding protein, TEVcs is the TEV peptide cleavage site (ENLYFQS), FKBP is FK506-binding protein, and EGFP is enhanced green fluorescent protein. The extent of proteolysis was evaluated by monitoring the accumulation of cleaved product via sodium dodecyl sulfate-polyacrylamide gel electrophoresis (SDS-PAGE) (Figure S9). Two designs, hyperTEV56 and hyperTEV60, exhibited significantly higher rates of cleavage of protein substrate compared to the parent TEVd, yielding 50% cleaved product at ∼4 hours of incubation while TEVd required 24 hours to reach an equivalent yield. The designs also outperformed other published TEV variants, with 30% turnover for superTEV, 15% turnover for TEV1Δ, and 50% turnover for S219V at 24 hours of incubation (Figures 3C and S10A). Straight-line fit of product accumulation and substrate depletion reveal catalytic efficiencies that corroborate those determined in the peptide assay (Figure S10B). The gains in catalytic efficiency are primarily due to increases in *k*_cat_, which could reflect a higher fraction of enzyme in a catalytically competent state (see below).

Analysis by CD spectroscopy of TEVd and the most active design, hyperTEV60, indicated an approximate melting temperature of 84 °C for hyperTEV60, 40 °C higher than that of TEVd (Figures 3D and S11), and to our knowledge, higher than any previously described TEV variant. To further probe stability of the designed variant, TEVd and hyperTEV60 were incubated at 30 °C for various times and then used in the peptide-coumarin cleavage assay. After 4 hours of incubation, hyperTEV60 retained 90% of its original cleavage activity while TEVd was reduced to 15% of its original activity (Figure 3E), indicating a significant improvement in benchtop stability.

Given that catalytic and substrate-binding residues were kept fixed during design with ProteinMPNN, it is notable that significant improvements in *k*_cat_ were observed with both the peptide and protein substrates. Mutations distal to the active site can influence catalytic activity through stabilization of catalytically productive conformational states^38,39^ or global conformational changes^40^. To investigate if stabilization of functional conformational states may be involved in activity enhancement, we performed microsecond molecular dynamics (MD) simulations on TEV-peptide complexes to probe the impact of the introduced mutations on overall protein dynamics. A general rigidification of loop regions distributed across the structure was observed in designs as compared to TEVd (Figure S12A). This backbone rigidification in distal regions not directly involved in substrate binding may be related to allosteric improvement of substrate binding as reflected by the 2- to 3-fold lower *K*_m_ values measured for the designed variants (Table 1). Rigidification in the region spanning residues 115 to 124 appeared to correlate with activity; the highest activity design, hyperTEV60, was most rigid, while TEVd and a design with no activity on the peptide substrate were most flexible in this region (Figure S12B). These trends were also observed in per-residue pLDDT analysis of AlphaFold2 ensemble predictions (Figure S12C). In all designs, we observed a decrease in the population of catalytically competent conformations of the Cys-His dyad (d_N-SH_) compared to TEVd, but this shift was least significant in hyperTEV60, in agreement with its higher relative *k*_cat_ (Figure S13). These notable differences may begin to explain how ProteinMPNN enables substantial activity enhancements without explicit design elements to improve function. It is also possible that the major contribution to the increase in *k*_cat_ is from an increase in the fraction of the protein in the catalytically competent state more globally.

## Conclusion

We show that the expression, stability, and function of native proteins can be improved using ProteinMPNN guided by available sequence and structural information. For both TEV protease and human myoglobin, multiple variants were identified which showed higher soluble yield and thermostability than the native protein starting point. The best of the TEV protease designs have higher apparent catalytic efficiency on peptide and protein substrates than the parent enzyme and previously reported variants. While the optimal number of residues to conserve to maintain (and perhaps enhance) function may have to be determined empirically for each case, the simplicity of our procedure and the compute efficiency and ease of use of ProteinMPNN make this straightforward, and the number of variants that need to be tested is far smaller than in typical experimental screens. We expect that our approach should be widely useful for improving the expression, stability, and function of biotechnologically important proteins.

## Acknowledgements

We thank A. Lauko, C. Norn, A. Roy, L. Stewart for helpful discussions. We thank L. Goldschmidt and K. VanWormer for computational and experimental support, respectively. We thank X. Li and M. Lamb for analytical services. We also thank Agencia Estatal Investigacion of Spain (PID2021-125946OB-I00, CEX2021-001136-S, predoctoral fellowship; G.J.O., R.N.F.) for support of this work. Crystallographic data were collected at The Advanced Light Source (ALS), which is supported by the director, Office of Science, Office of 20 Basic Energy Sciences, and US Department of Energy under contract number DE-AC02-05CH11231. Funding was also provided by a National Science Foundation (NSF) grant CHE-1629214 (A.K.B.), the Air Force Office of Scientific Research (S.J.P.), a Defense Threat Reduction Agency grant HDTRA1-19-1-0003 (S.J.P.), the Bill and Melinda Gates Foundation grant #OPP1156262 (A.K., J.C.), the National Institute of Health’s National Institute of Allergy and Infectious Disease (R0AI160052, A.B.K.), the DARPA program Harnessing Enzymatic Activity for Lifesaving Remedies (HEALR) (HR0011-21-2-0012, A.K.B.), the Audacious Project at the Institute for Protein Design (L.F.M., A.K., J.C., E.B., A.K.B., D.B.), the Howard Hughes Medical institute (I.K., Y.K., D.B.), the Open Philanthropy Project Improving Protein Design Fund (K.H.S., S.J.P., I.K., B.I.M.W., J.D., J.W., Y.K., A.K., J.C., E.B., A.K.B., D.B.), Schmidt Futures (J.W.), a grant from the National Science Foundation (NSF) (DBI 1937533; D.B.), the Department of Energy ARPA-E Grant #2459-1671 (D.B.), an EMBO long-term fellowship (ALTF 139-2018; B.I.M.W.), a Washington Research Foundation Fellowship (S.J.P., J.W.), an Alfred P. Sloan Foundation Matter-to-Life Program Grant (G-2021-16899; D.B.), an EMBO Non-Stipendiary Fellowship (ALTF 1047-2019; L.F.M.), a Human Frontier Science Program Cross-Disciplinary Fellowship (LT000838/2018-C (I.K.), LT000395/2020-C (L.F.M.)), the National Institute of Health (R35 GM124773; N.J.) and a gift from Microsoft (J.D., D.B.).

## Methods

### Fixed residue selection for TEV protease

Active site positions were defined as residues containing backbone atoms within 7 Å of the substrate or sidechain atoms within 6 Å of the substrate in the ligand-bound crystal structure of autolysis resistant S219D (PDB: 1LVM). For enzyme targets, highly conserved residues were also fixed during sequence redesign. Highly conserved residues were determined with multiple sequence alignments (MSA). To generate the MSA, four iterative HHblits searches^41^ were performed against the UniRef30 database (accessed June 30, 2020) at E-value cutoffs of 1e-50, 1e-30, 1e-10, and 1e-4, and the final result was filtered for 90% identity redundancy, 50% coverage, and 30% minimum query identity. Within the sequence alignment, we identified the frequency of each amino acid at each position and found the most highly conserved amino acid identity at each position. We then ranked each position by how highly conserved the most frequent amino acid identity was, and selected the top 30%, 50%, and 70% most conserved positions to fix during sequence design.

### ProteinMPNN design of myoglobin

For fixed-backbone sequence redesign, the crystal structure of human myoglobin (PDB: 3RGK) was used as input to ProteinMPNN, and 17 positions located around the heme were excluded from design. Three temperatures (0.1, 0.2, and 0.3) were sampled, with 20 sequences generated per temperature. Cysteine and methionine were excluded from the amino acid identities that could be installed during design. A model of ProteinMPNN trained with 0.2 Å noise applied to training set protein backbones was used to perform sequence generation. For combined sequence and backbone redesign, two strategies were employed. First, the sequence and structure from the crystal structure of human myoglobin (PDB: 3RGK) were input to RF_joint_, with the N- and C-termini and loop region between helices 5 and 6 masked, to generate new secondary structure in these regions (RoseTTAFold joint inpainting). Ten backbones were generated with this strategy. In a more aggressive strategy, helix 4 and its adjoining loops, as well as both termini and the loop joining helices 5 and 6, were masked. Twenty backbones were generated with this strategy. Following backbone redesign, 60 sequences were generated per backbone with ProteinMPNN, keeping heme-binding positions fixed as described above.

Sequences generated with ProteinMPNN were predicted with AlphaFold2, using model 4 with 10 recycling steps. Structural templating with MSAs was not used for prediction. Designs with only sequence redesign were filtered to Cα RMSD < 1.0 Å and pLDDT > 85.0. Designs with sequence redesign and backbone redesign on the termini and the loop connecting helices 5 and 6 were filtered to Cα RMSD < 0.8 Å and pLDDT > 90.0. Designs with sequence redesign and backbone redesign on the termini, helix 4, and the loop connecting helices 5 and 6 were filtered to Cα RMSD < 0.6 Å and pLDDT > 90.0 (see Figure S2 for details). Predicted models passing these criteria were finally evaluated by eye and those recapitulating finer structural details of the heme binding pocket (low backbone deviation after global alignment to the structure of 3RGK; close agreement with the placement of heme-coordinating histidine side chain) were selected for experimental testing. Four designs generated with only sequence design, and 16 designs with sequence and backbone design were selected for experimental testing (10 with both loops remodeled, and 6 with one loop).

### ProteinMPNN design of TEV protease

The crystal structure of TEVd (PDB: 1LVM) was used as structural input to ProteinMPNN, and active site and conserved residues were excluded from design. Cysteine was excluded from the amino acid identities that could be installed during design. Three temperatures (0.1, 0.2, and 0.3) were sampled during design. A model of ProteinMPNN trained with 0.2 Å noise applied to training set protein backbones was used to perform sequence generation. 24 sequences were generated with only the active site residues fixed, 24 sequences were generated with the active site and the 30% most highly conserved positions fixed, 48 sequences were generated with the active site and the 50% most highly conserved positions fixed, and 48 sequences were generated with the active site and the 70% most highly conserved positions fixed.

Sequences generated with ProteinMPNN were predicted with AlphaFold2, using model 3 with 6 recycling steps. Both designs and native TEV predicted with low confidence if given only the single sequence and minimal recycling steps. We found that structural templating with MSAs was necessary for accurate prediction. To generate MSAs of each design for structure prediction, the MSA of the parent sequence was used, and the parent sequence was swapped for the design sequence. All sequences generated were predicted with Cα RMSD < 2.0 Å and pLDDT > 85.0 and were predicted to maintain critical structural features in the active site. Thus, all were ordered for experimental characterization.

### Expression and purification of myoglobin designs

Double-stranded DNA fragments encoding the designs (codon-optimized for bacterial expression) were purchased from Integrated DNA Technologies (IDT) as eBlocks™ Gene Fragments. Following the Golden Gate cloning protocol,^14^ the DNA fragments encoding design sequences and including overhangs suitable for a *Bsa*I restriction digest were cloned into a custom pET29b(+) target vector containing lethal ccdb gene, and C-terminal SNAC^42^ and hexahistidine tags (#191551, Addgene). This yielded final expressed sequences as: MSG<design>GSGSHHWGSTHHHHHH. Assembled plasmids containing the designs were transformed into *E. coli* BL21(DE3) by heat shock. DNA was incubated on ice with competent cells for 30 minutes, followed by 30 second heat shock at 42 °C, and 2 minute incubation on ice. 100 μL rich medium (super optimal broth with catabolite repression) was added to transformed cells and samples were incubated at 37 °C, 1050 r.p.m. on a Heidolph shaker for 1 hour. The cells were subsequently spread on LB-agar plates containing 100 μg/mL kanamycin and incubated at 37 °C under 220 r.p.m. shaking for 18 hours. Single colonies were picked, and the DNA fragments encoding the designs were amplified following a colonyPCR protocol using GoTaq® Green DNA polymerase master mix (#M7122; Promega) and T7 reverse and forward primers. The PCR products identified to contain DNA of appropriate size (∼600 bp) based on agarose gel (1.2%) electrophoresis with SybrSafe dye were sent to Sanger sequencing (GeneWiz/Azenta) for sequence-verification. Single colonies containing the correct design sequences were grown up in 5 mL TB-II media containing 50 μg/mL kanamycin, over 16 hours at 37 °C. 2 mL of the grown culture was used to inoculate 40 mL TB-II media containing 50 μg/mL kanamycin and the rest used for plasmid extraction following the Qiagen QIAprep MiniPrep protocol. The 40 mL cultures were grown at 37 °C for 4 hours, after which protein expression was induced with the addition of 1 mM IPTG, and the cultures were incubated at 18 °C for 20 hours. Pellets were harvested by centrifugation at 4,198 g for 8 minutes and resuspended in a lysis buffer containing 25 mM Tris-HCl, 300 mM NaCl, 25 mM imidazole, 0.01 mg/mL DNAse, 0.1 mg/mL lysozyme, and a Pierce protease inhibitor tablet. Lysis was performed by ultrasonication (13 mm probe, 2.5 mins, 10s on, 10s off, 65% amplitude). Lysate was collected by centrifugation at 15,000 xg for 20 minutes and applied to Ni-NTA resin that was equilibrated with wash buffer (25 mM Tris-HCl, 300 mM NaCl, 25 mM imidazole, pH 8.0). The resin was washed with 50 column volumes (CV) of wash buffer. Protein was eluted with 1.2 CV of elution buffer (25 mM Tris-HCl, 300 mM NaCl, 300 mM imidazole, pH 8.0) and further purified via size exclusion chromatography (SEC) using a Superdex Increase 75 10/300 GL column (GE Healthcare) on ÄKTAxpress (GE Healthcare) instrument at 0.8 mL min-1 flow rate. The monomeric or smallest oligomeric fractions of each run (eluting at approximately 15 ml) were collected. The obtained chromatograms are presented in Figure S6.

Yields of purified hemoproteins were determined based on the absorbance of the Soret maximum (407-413 nm). The corresponding extinction coefficients were measured for each protein using the hemochromagen assay, according to the method of Berry and Trumpower.^43^ A reported extinction coefficient of 188 mM^-1^·cm^-1^ was used for native myoglobin.^44^

### Expression and purification of TEV designs

Double-stranded DNA fragments encoding the designs (codon-optimized for bacterial expression) were purchased from Integrated DNA Technologies (IDT) as eBlocks™ Gene Fragments. Following the Golden Gate cloning protocol,^14^ the DNA fragments encoding design sequences and including overhangs suitable for a *Bsa*I restriction digest were cloned into a custom pET29b(+) target vector containing lethal ccdb gene, and C-terminal SNAC^42^ and hexahistidine tags (#191551, Addgene). This yielded final expressed sequences as: MSHHHHHHSG<design>GS. Vectors containing TEV designs were transformed into *E. coli* BL21(DE3) by heat shock. DNA was incubated on ice with competent cells for 30 minutes, followed by 10 second heat shock at 42 °C, and 2 minute incubation on ice. 100 μL rich medium (super optimal broth with catabolite repression) was added to transformed cells and samples were incubated at 37 °C, 1050 rpm on a Heidolph shaker for 1 hour. Entire transformations were transferred to 900 μL of TBM-5052 autoinduction expression medium containing 50 μg/mL Kanamycin. Expression cultures were incubated at 37 °C, 1050 rpm for 20 hours. Pellets were harvested by centrifugation at 4,000 g for 10 minutes and lysed with BPER lysis reagent containing 6.25 Units/mL benzonase (4 uL / 40 mL at 250 U/μL), 0.1 mg/mL lysozyme, and 1 mM PMSF. Lysate was collected by centrifugation at 4,000 xg for 20 minutes and applied to Ni-NTA resin that was equilibrated with wash buffer (20 mM Tris-HCl, 300 mM NaCl, 25 mM imidazole, pH 8.0). The resin was washed with 25 column volumes (CV) of wash buffer. Protein was eluted with 250 μL of elution buffer (20 mM Tris-HCl, 300 mM NaCl, 540 mM imidazole, pH 8.0) and further purified via size exclusion chromatography (SEC) in an S75 5/150 GL increase column (GE Healthcare). Protein collected from SEC was normalized to 1 μM where possible.

In scale-up experiments, 50-mL cultures of TBM-5052 autoinduction media with 50 μg/mL Kanamycin were inoculated with a scrape of transformed competent cells from glycerol stock and grown at 37 °C, 200 rpm for 20 hours. Cells were harvested by centrifugation at 10,000 xg for 10 minutes, resuspended in 30 mL of wash buffer (20 mM Tris-HCl, 300 mM NaCl, 25 mM imidazole, pH 8.0) containing 0.01 mg/mL DNase, 0.1 mg/mL lysozyme, and a protease inhibitor tablet (Thermo Scientific Pierce), and lysed by sonication. Lysate was collected via centrifugation at 18,000 xg for 40 minutes and applied to Ni-NTA resin that was equilibrated with wash buffer. The resin was washed with 30 CV of wash buffer. Protein was eluted with 4 mL of elution buffer and concentrated to 1 mL in a 3K protein concentrator (Millipore Sigma). Concentrated protein was purified by SEC as described above.

### Expression and purification of MBP-TEVcs-FKBP-EGFP construct

The protease substrate FKBP-EGFP was cloned into an *E. coli* expression vector containing an N-terminal maltose binding protein (MBP), a TEV protease recognition site, and a C-terminal His-6 tag. The FKBP-EGFP coding sequence was obtained from Addgene #106924, with a 4X GGS linker between FKBP and EGFP. Vector containing the protease substrate was transformed into *E. coli* BL21(DE3) by heat shock. Cells were transferred to 4 0.5-L LB medium cultures with 50 μg/mL Kanamycin and incubated at 37 °C, 200 rpm until optical density reached 0.5 AU, at which point expression was induced with 1 mM IPTG. Temperature was reduced to 18 °C and cells were incubated for an additional 18 hours. Cells were harvested by centrifugation at 10,000 xg for 10 minutes, resuspended in 30 mL of wash buffer (20 mM Tris-HCl, 300 mM NaCl, 25 mM imidazole, pH 8.0) containing 0.01 mg/mL DNase, 0.1 mg/mL lysozyme, and a protease inhibitor tablet (Thermo Scientific Pierce), and lysed by sonication. Lysate was collected via centrifugation at 18,000 xg for 40 minutes and applied to Ni-NTA resin that was equilibrated with wash buffer. The resin was washed with 30 CV of wash buffer. Protein was eluted with elution buffer until resin no longer appeared yellow and concentrated to 1 mL in a 3K protein concentrator (Millipore Sigma). Concentrated protein was purified by SEC as described above.

### Kinetic characterization of designed proteases

Designs were initially screened for activity on a peptide-coumarin conjugate substrate (WuXi) of the TEV recognition sequence (ENLYFQ) fused to a fluorescent coumarin derivative, 7-amino-4-trifluoromethylcoumarin. The N-terminus of the peptide bears an acetyl modification and the C-terminus is conjugated to the coumarin group via an amide bond. Initial activity screen was performed in 50 mM Tris-HCl, 50 mM NaCl, pH 8.0 buffer containing Freshly prepared 2 mM DTT. Reactions contained 500 nM protein and 10 μM substrate at a total volume of 30 μL. Protein and substrate were rapidly mixed and monitored for fluorescence at excitation 400 nm, emission 492 nm at room temperature (RT) for 5 hours in a BioTek Synergy Neo2 microplate reader.

For detailed kinetic characterization, reactions were performed in 50 mM Tris-HCl pH 8.0 containing 50 mM NaCl, 1mM EDTA, and freshly prepared 2 mM DTT. For TEV redesigns, reactions contained 50 nM protein and substrate concentration ranging from 0.1 μM to 10 μM at a total volume of 30 μL. Protein and substrate were rapidly mixed and monitored for fluorescence at excitation 400 nm, emission 492 nm at RT for 2 hours in a BioTek Synergy Neo2 microplate reader. Fluorescent signal was converted to concentration of cleaved coumarin product using a calibration curve of 7-amino-4-trifluoromethylcoumarin. Reactions were performed in triplicate and each technical replicate was separately fitted to a Michaelis Menten model. Expressed uncertainty in *k*_cat_ and *K*_m_ is the standard deviation between technical replicates.

### Screening of designed proteases on fusion protein MBP-TEVcs-FKBP-EGFP

Reactions were performed in 50 mM Tris-HCl, 50 mM NaCl, 1mM EDTA, pH 8.0 buffer containing freshly prepared 2 mM DTT. Reactions contained 60 nM protein and substrate concentrations ranging from 2 μM to 17 μM. Reactions were incubated at 30 °C and at 0, 1, 2, 4, 8, and 24 hours, 10 μL aliquots were quenched in 10 μL of 2X Laemmli loading buffer and subsequently frozen in liquid nitrogen. Samples were analyzed by SDS-PAGE and imaged for EGFP fluorescence at 488 nm on a LI-COR Odyssey M imager. Band intensities were quantified with ImageJ software and converted to concentration using a standard curve prepared of known amounts of cleaved substrate with fluorescence gel imaging. A straight-line fit was applied to the initial velocities using GraphPad Prism. Points represent the averages of 3 technical replicates and error bars represent the standard deviations.

### Benchtop stability characterization of TEV redesigns

Samples of purified enzyme were incubated at 30 °C for 0.5, 1, 2, 4, 8, 18, or 24 hours before being used in the previously described peptide-coumarin cleavage assay. Activity of samples was defined as initial rate of turnover and normalized to initial rate at incubation of t = 0 hrs.

### Spectrophotometric measurementsUV

Vis spectra of purified holo-proteins (myoglobin variants) in the 230-800 nm range were collected using the Jasco Spec V750 spectrophotometer and 10 mm pathlength cuvette. To observe changes in the spectral properties of bound heme at increasing temperatures, UV−Vis spectra were collected at every 10 °C intervals between 25 °C and 95 °C. Temperature was increased at the rate of 5 °C min-1, and spectra were acquired after the temperature had stabilized to within 0.5 °C of target temperature for 5 seconds. Measurements were performed with 20 μM solutions of purified holoprotein in TBS buffer (25 mM Tris-HCl, 300 mM NaCl, pH 8).

### Circular dichroism spectroscopy

To determine secondary structure and thermostability of the designs, far-ultraviolet circular dichroism (CD) measurements were carried out on a JASCO J-1500 instrument using a 1 mm pathlength cuvette. Samples of purified protein were prepared at 1.0 mg/mL in 20 mM sodium phosphate, 50 mM potassium fluoride, pH 8.0 (TEV protease) or at 0.4 mg/mL in 25 mM Tris, 20 mM NaCl, pH 8.0 (myoglobin). The temperature of the sample was scanned from 25 °C to 95 °C with full spectrum scans from 190 nm to 260 nm performed after each 10 degree increment. The signal at 216 nm was plotted over the temperature gradient and fitted to a Boltzmann sigmoidal curve with GraphPad Prism 9. *T*_m_ values were calculated from the inflection point.

### Mass spectrometry analysis

MS data for dnHEM1 variants were acquired on an Agilent 1200series LC G6230B TOF LC-MS with an AdvanceBio RP-Desalting column (A: H2O with 0.1% Formic Acid, B: Acetonitrile with 0.1% Formic Acid). The final protein concentrations were adjusted to 1-2 mg/mL in 25 mM Tris-HCl, 300 mM NaCl, pH 8.2. Subsequent data deconvolution was performed in Bioconfirm using a total entropy algorithm. All data are presented in Supplementary Table 1.

### Molecular dynamics simulations

Structures generated with AlphaFold2^15^ were used as starting geometries. For the protein-substrate complexes, substrate peptide was superimposed onto AlphaFold2 structures using the crystallographic structure of catalytically active TEV protease (PDB: 1LVM) as a template. Simulations were carried out with AMBER 20^45^ implemented with the ff14SB force field for the protein and substrate peptide, and the general Amber force field (GAFF2)^46^ for the substrate peptide C-terminal fluorescent probe (7-amino-4-(trifluoromethyl)coumarin). Parameters were generated with the *antechamber* module of AMBER, combining ff14SB and GAFF2 force fields and with partial charges set to fit the electrostatic potential generated with HF/6-31G(d) using the RESP method.^47^ The charges were calculated according to the Merz-Singh-Kollman scheme using Gaussian 16.^48^ Binding-site histidine residue (H46) was modeled in its Nδ1-H tautomeric state (corresponding to residue name HID in Amber). Initial structures were neutralized with either Na^+^ or Cl^−^ ions and set at the center of a cubic TIP3P^49^ water box with a buffering distance between solute and box of 10 Å.

A two-stage geometry optimization approach was performed. The first stage minimizes only the positions of solvent molecules and ions, and the second stage is an unrestrained minimization of all the atoms in the simulation cell. The system was then heated by incrementing the temperature from 0 to 300 K under a constant pressure of 1 atm and periodic boundary conditions (PBC). Harmonic restraints of 10 kcal/mol were applied to the solute, and the Andersen temperature coupling scheme^50,51^ was used to control and equalize the temperature. The time step was kept at 1 fs during the heating stages, allowing potential inhomogeneities to self-adjust. Water molecules were treated with the SHAKE algorithm^52^ such that the angle between the hydrogen atoms is kept fixed through the simulations. Long-range electrostatic effects were modeled using the particle mesh Ewald method.^53^ An 8 Å cut-off was applied to Lennard-Jones interactions. The system was equilibrated for 2 ns with a 2 fs time step at a constant volume and temperature of 300 K. Ten independent production trajectories were then run for additional 1000 ns under the same simulation conditions, leading to accumulated simulation times of 10 μs for each system. Root mean square (rms) fluctuations and interatomic distance analyses were carried out with the *cpptraj* module of AMBER.

## Data availability

The atomic coordinates and structure factors of the reported structure have been deposited in the Protein Data Bank under accession code 8U5A for dnMb19.

## Author contributions

K.H.S., R.N.F., and I.K. performed experiments and analyzed data. K.H.S., R.N.F., I.K., S.J.P., B.I.M.W., L.F.M., and D.B. planned experiments. A.K., J.C., B.S., and A.K.B. collected and processed data for X-ray crystallography. N.J. provided materials and expertise. K.H.S., R.N.F., I.K., S.J.P., B.I.M.W., L.F.M., J.D., J.W., Y.K., and D.B. developed the design protocol. K.H.S., I.K., S.J.P, and D.B. wrote and edited the paper. D.B. and G.J.O. offered supervision throughout the project. All authors read and contributed to the manuscript.

