## Supplementary Data for "Improving protein expression, stability, and function with ProteinMPNN"

#### Content

1. Supplementary Figures 1
2. Crystallographic data 17
3. Sequence information 19
4. Computational details 29
5. References 32

#### 1. Supplementary Figures

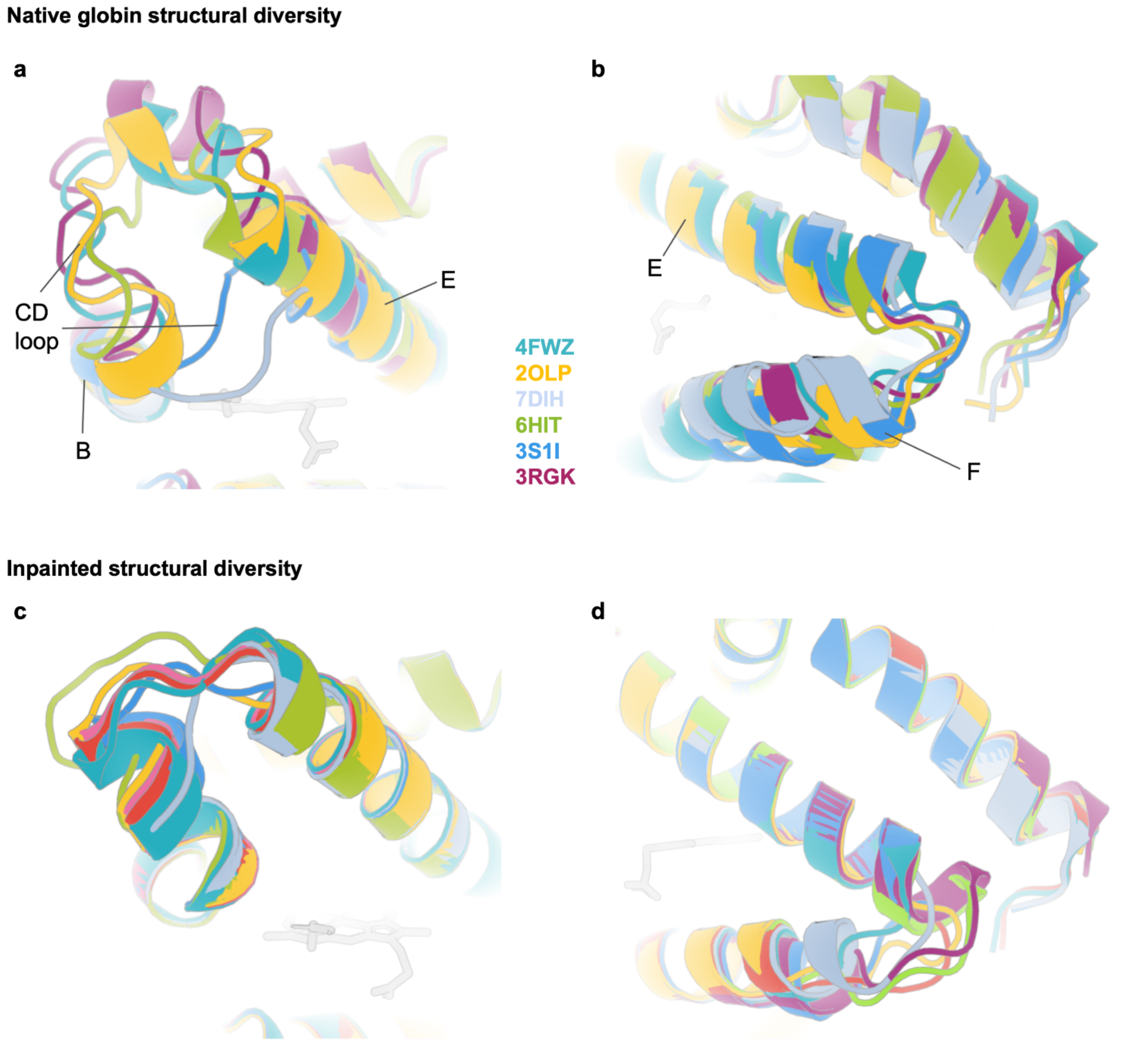


**Figure S1.** Inpainting samples different backbone structures compared to native globins. (A) Diversity of the CD-loop region in selected native globins. (B) Diversity in the loop connecting helices E and F in selected native globins. (C) Diversity of inpainted motifs replacing the CD-loop region. (D) Diversity of inpainted loops connecting helices E and F.

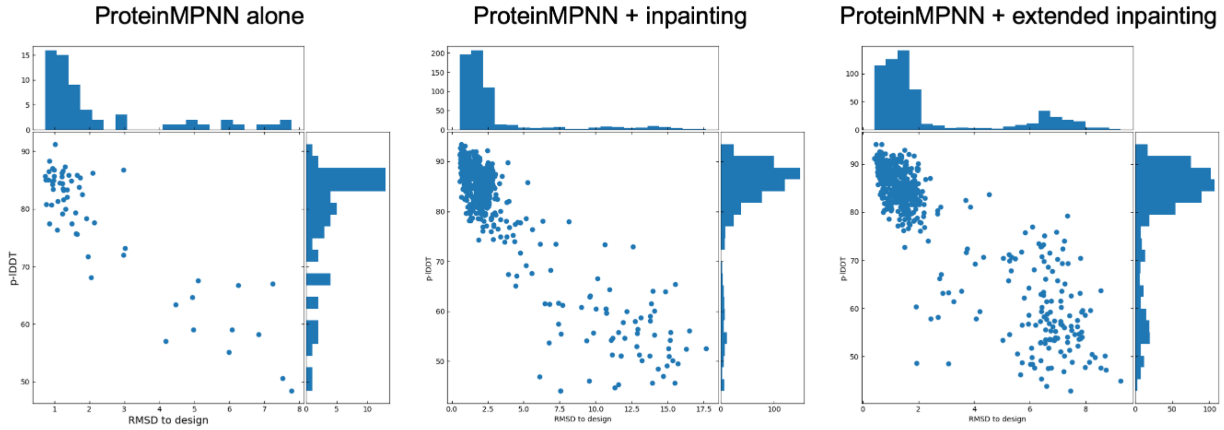


**Figure S2.** Extensive backbone remodeling with RoseTTAFold joint inpainting improves structure prediction metrics. Designs made with only sequence redesign had the lowest-scoring structure prediction metrics (pLDDT and RMSD to design model) amongst all designs, while designs subjected to the most aggressive backbone remodeling strategy scored the highest in these metrics.

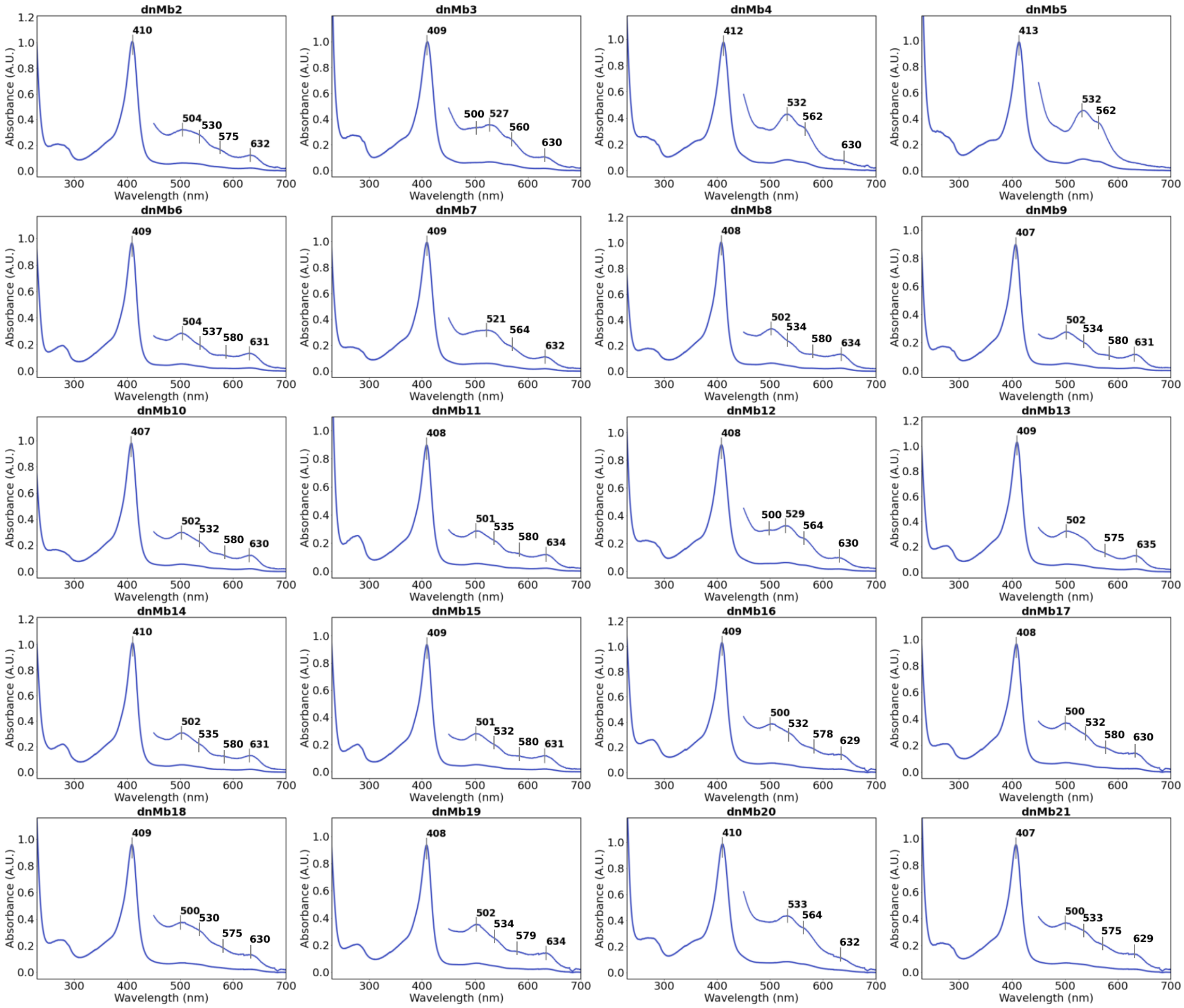


**Figure S3.** UV/Vis spectra of myoglobin variants. Spectroscopic data of most designs is in close agreement with that of native myoglobin (Soret maximum at 409 nm; Q band features at 500, 537, 582 and 630 nm), suggesting pentacoordinate heme-binding. A few designs (3, 4, 5, 12, 20) show some degree of hexacoordinate heme-binding (potentially through incorporation of imidazole from the purification buffer), indicated by the major Q band features at ~530 and ~560 nm. Spectra were recorded in a buffer containing 25 mM Tris-HCl and 300 mM NaCl at pH 8.2.

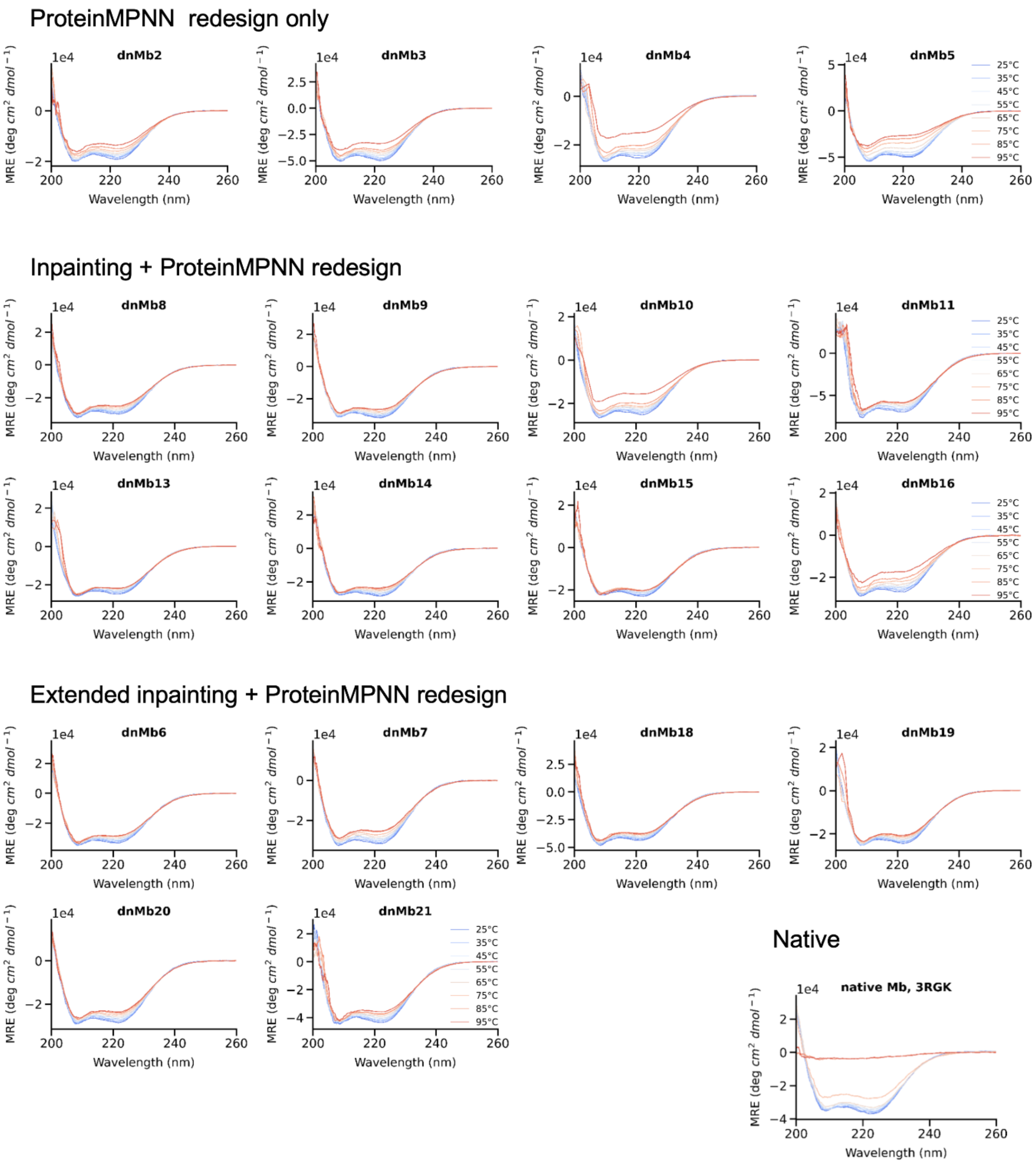


**Figure S4.** Many myoglobin designs show increased thermostability over parent. CD spectroscopy signal of myoglobin designs and parent sequence nMb over a temperature gradient from 25 °C to 95 °C indicates elevated resistance to unfolding in designs. CD signal reported in molar residue ellipticity (MRE).

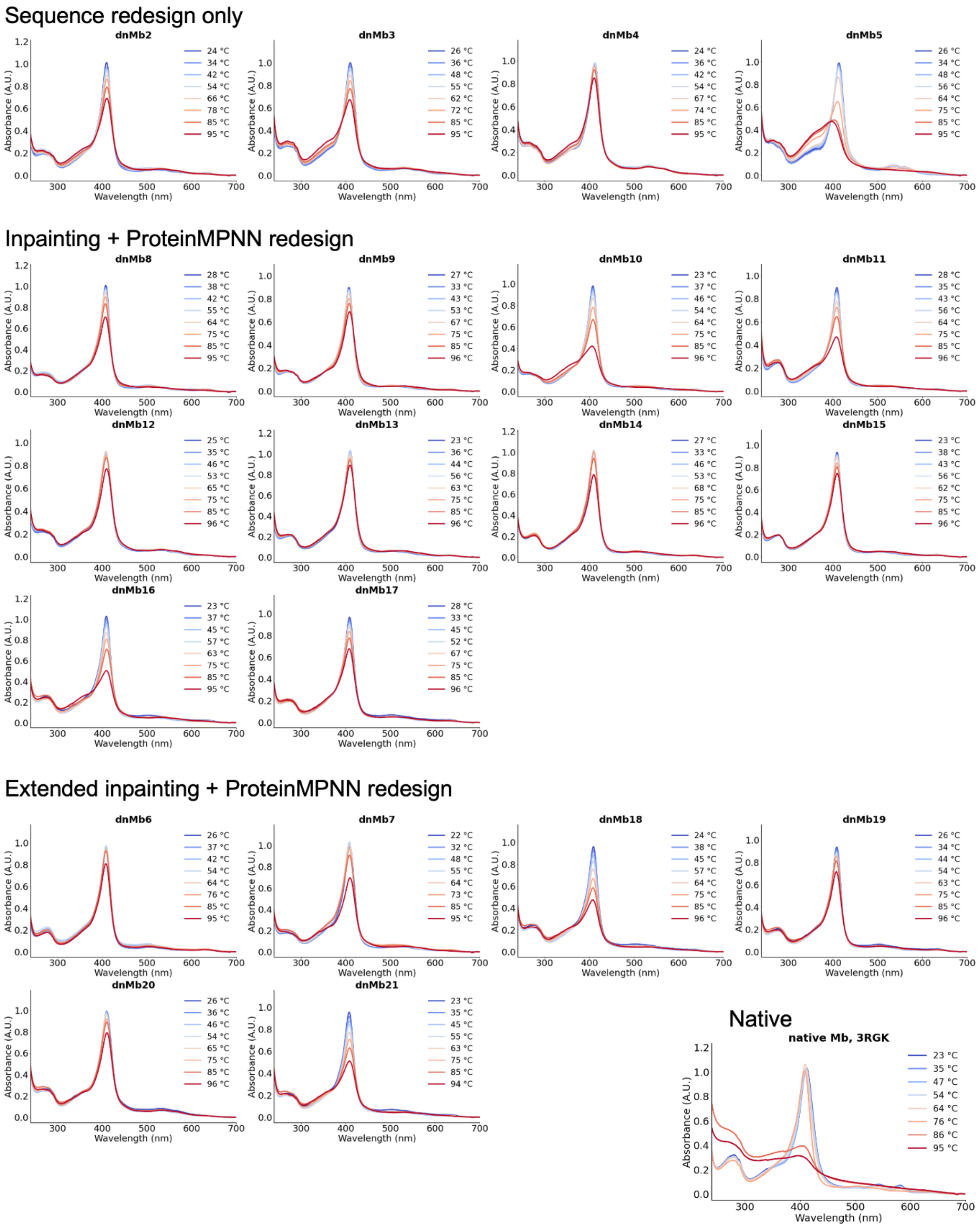


**Figure S5.** Myoglobin designs retain heme binding at higher temperatures than parent. Heme binding as measured by UV/Vis absorbance over a temperature gradient from 25 °C to 95 °C indicates retention of function at higher temperatures in designs. Higher melting temperatures of designs indicate more temperature-stable binding sites.

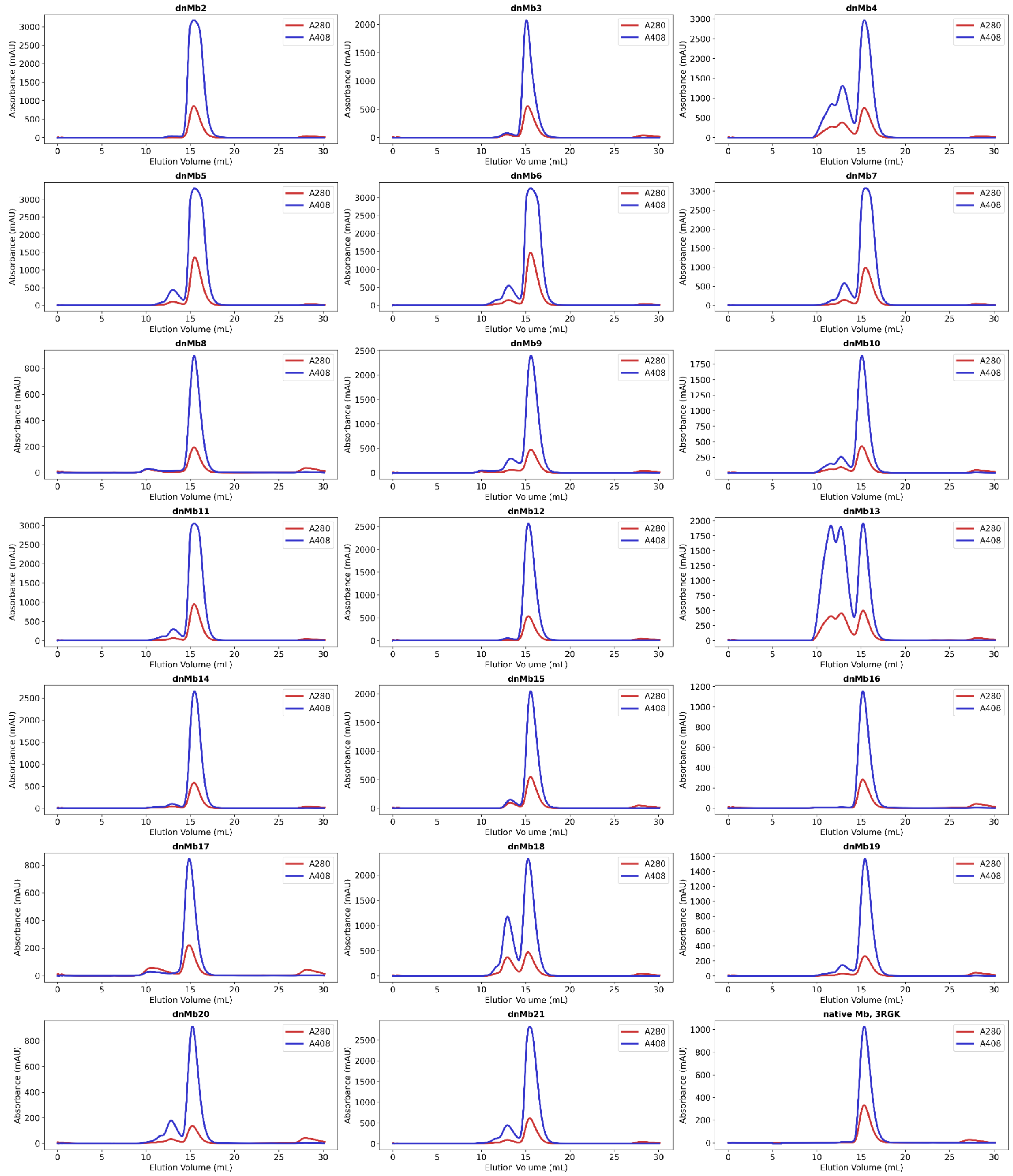


**Figure S6.** Size-exclusion chromatograms of heme-loaded myoglobin variants. Data were collected using a Superdex Increase 75 10/300 GL column (GE Healthcare) in a buffer containing 25 mM Tris-HCl and 300 mM NaCl at pH 8.2. Void volume of the column is 8.5 mL. Blue chromatograms were obtained by following the absorbance at 408 nm, indicating elution of heme-containing species. Red chromatograms were obtained from absorbance at 280 nm.

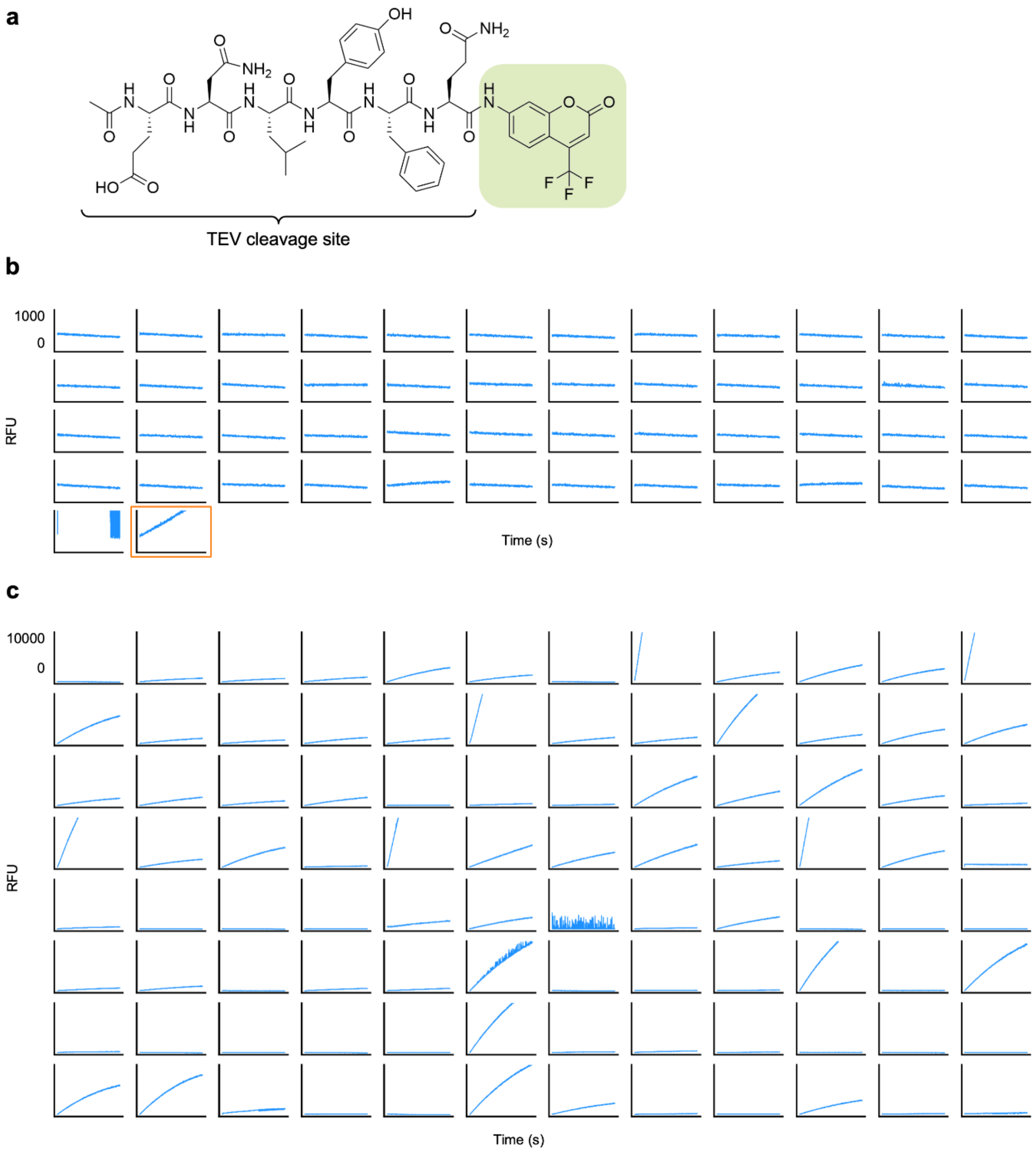


**Figure S7.** Initial screen of proteolytic activity on fluorescent reporter substrate. Pure protein was normalized to 1 µM and assayed against a single concentration of substrate, AFC, in an initial screen for catalytic turnover. (A) Structure of the peptide-coumarin substrate, AFC, used to assay proteolytic activity. (B) Raw fluorescence data (in raw fluorescence units, RFU) for designs generated with only active site residues fixed or with active site residues and 30% most conserved residues fixed during design. TEVd plot outlined in orange. (C) Raw fluorescence data for designs generated with active site residues fixed and 50% most conserved residues fixed or with active site residues fixed and 70% most conserved residues fixed.

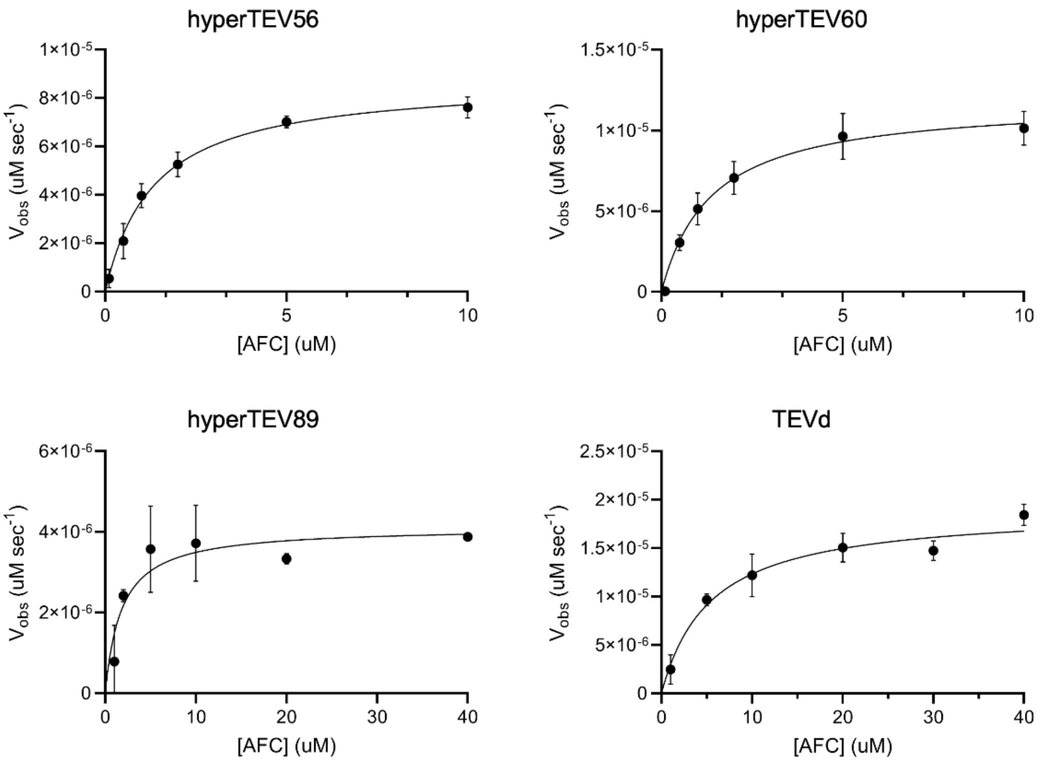


**Figure S8.** Michaelis Menten kinetics of TEV redesigns and parent. Michaelis Menten plots for three TEV designs and TEVd. Error bars represent standard deviation from three technical replicates.

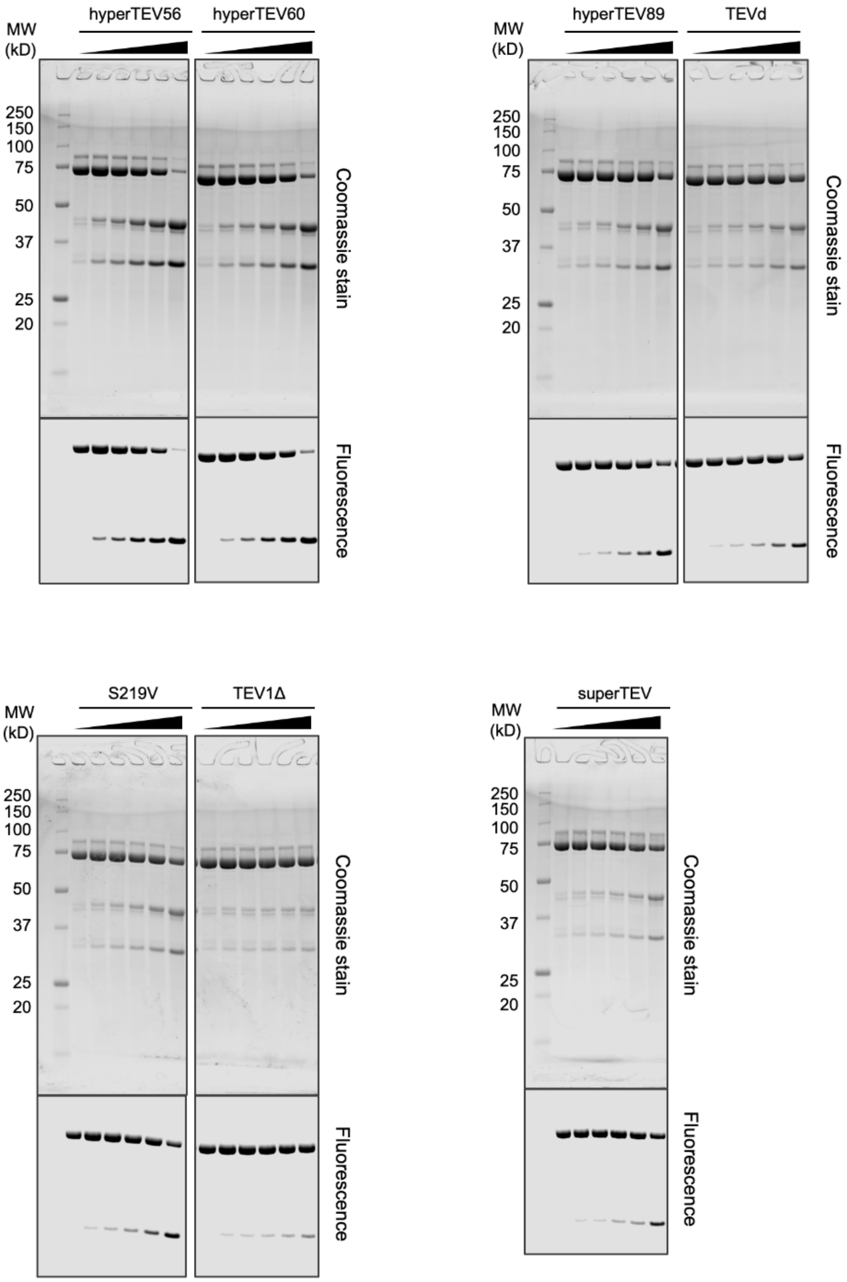


**Figure S9.** SDS-PAGE gels of protein substrate cleavage by TEV designs. Protein standard molecular weight ladder is shown on the left, with molecular weight markers indicated in kD. For each gel, the coomassie-stain is shown above and the EGFP fluorescence image is shown below.

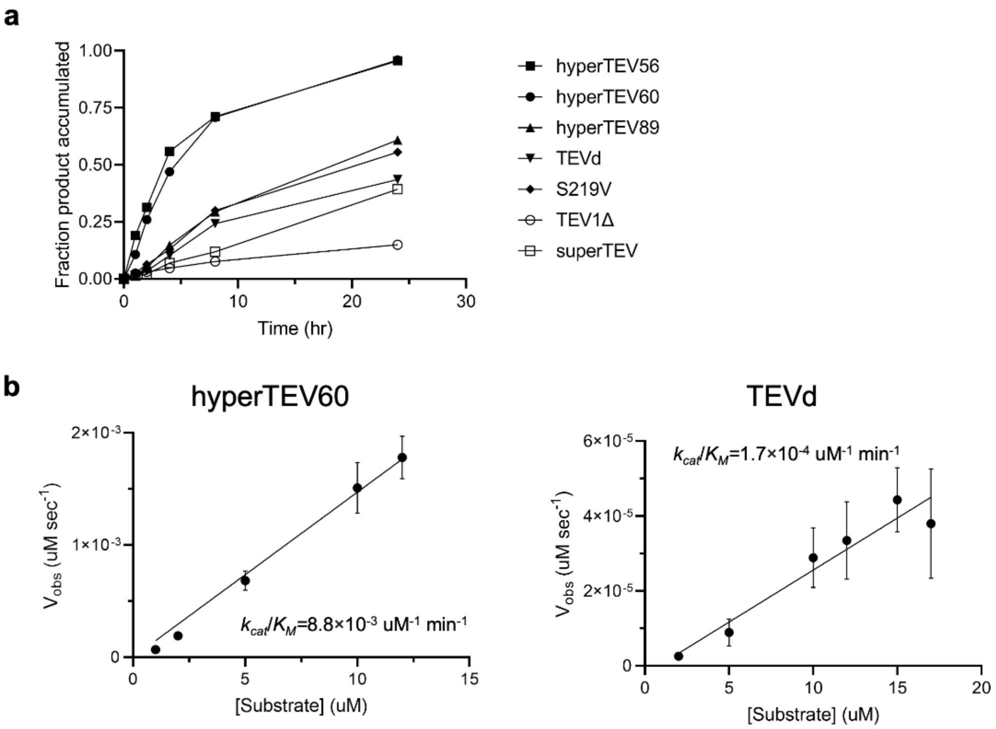


**Figure S10.** Activity of TEV redesigns in gel-based activity assay. (A) Plot of accumulated product normalized to fluorescence intensity of uncleaved substrate over time. Fluorescence intensity was quantified with ImageJ software. Designs hyperTEV56 and hyperTEV60 show increased turnover rate compared to reported TEV variants. (B) Straight-line fit for initial turnover rates in gel assay for hyperTEV60 and TEVd. Curves were fitted from monitoring of substrate depletion for hyperTEV60 and production accumulation for TEVd. Error bars represent standard deviation from three technical replicates.

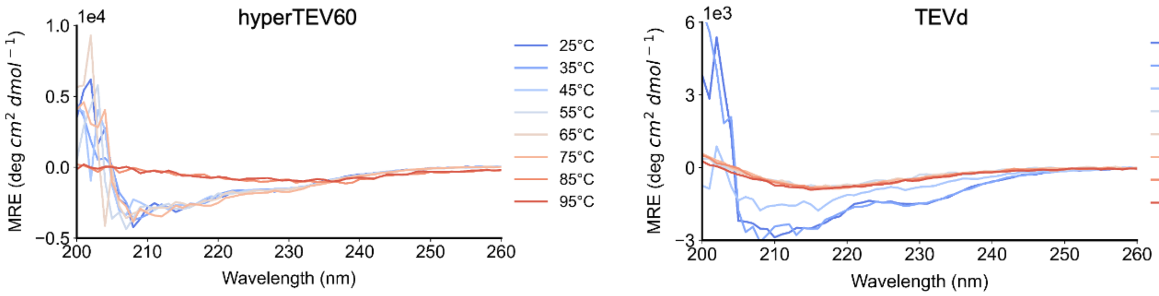


**Figure S11.** TEV design shows increased thermostability over parent. CD spectroscopy signal of hyperTEV60 and TEVd over a temperature gradient from 25 °C to 95 °C indicates elevated resistance to unfolding in ProteinMPNN design hyperTEV60. CD signal reported in molar residue ellipticity (MRE).

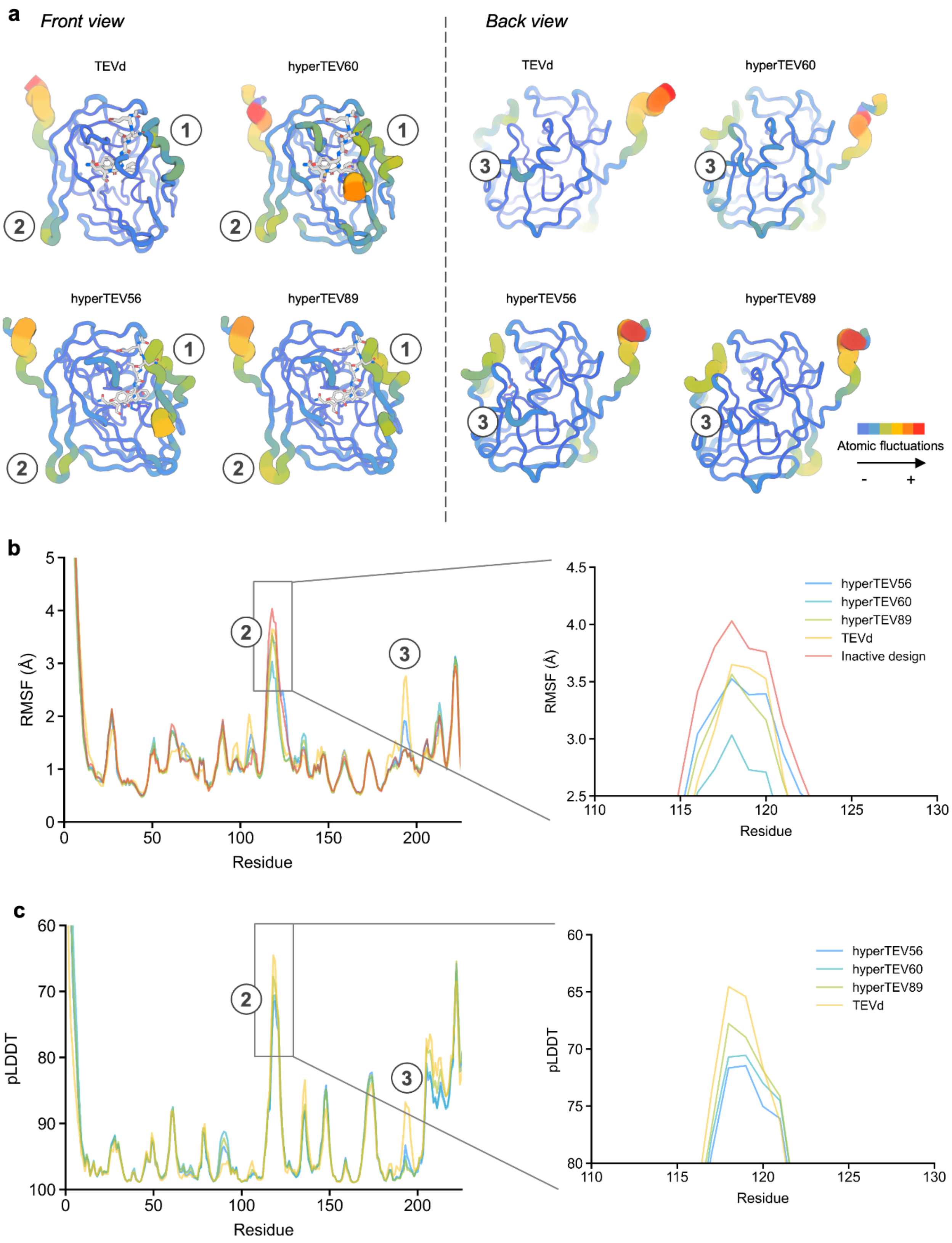


**Figure S12.** Designs show trends in rigidification and activity in molecular dynamics simulations. (A) Molecular dynamics (MD) simulations revealed trends of rigidification of several loops (marked with numbered circles) in the redesigned structures as compared to the parent. Directly adjacent to the peptide binding site, region 1 shows diminishing mobility in hyperTEV60 and other redesigns as compared to TEVd. An internal loop designated as region 3 shows significant loss of atomic fluctuation relative to TEVd. (B) Root-mean-square (RMSF) of designs in region 2 denoted in (A) shows a positive correlation between activity and rigidification, with TEVd and a design inactive on the peptide substrate showing most flexibility in this region. (C) Per-residue pLDDT values from AlphaFold2 ensemble prediction exhibit similar trends of increased rigidification in more active designs.

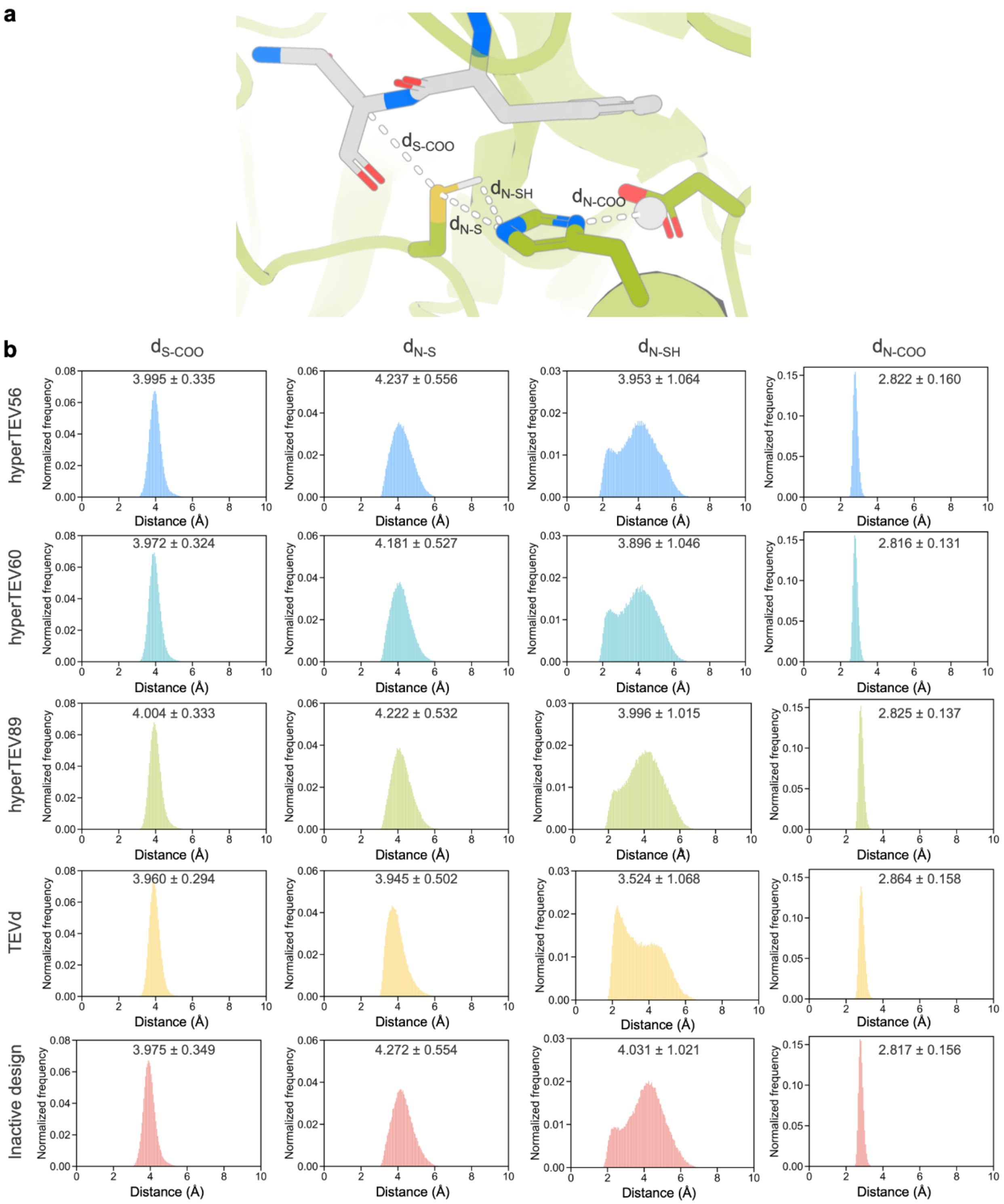


**Figure S13.** Population of catalytically competent dyad conformers correlates with activity in molecular dynamics simulations. (A) Key distances in the TEV catalytic triad as shown on TEVd. Peptide substrate is shown in gray. (B) Distances of key interactions in the catalytic triad were measured across MD simulations. Average distances for each interaction in each TEV variant are inset. Catalytically competent conformers of the Cys-His dyad ($d_{N\text{-}SH}$) are less populated in designs as compared to TEVd. Among designs, the highest activity variant hyperTEV60 has the highest percentage of competent dyad conformers.

**Table S1.** Myoglobin sequence similarity analysis against UniRef100

| Variant | Design method | Sequence similarity with parent (3RGK) | Highest sequence similarity | Most similar UniRef100 ID |
|---|---|---|---|---|
| dnMb2 | proteinMPNN only | 47% | 51% | P02182 |
| dnMb3 | proteinMPNN only | 49% | 52% | UPI00148EC9BB |
| dnMb4 | proteinMPNN only | 51% | 55% | P02182 |
| dnMb5 | proteinMPNN only | 46% | 51% | Q0KIY3 |
| dnMb6 | Inpaint CD+EH; MPNN | 40% | 44% | A0A8C5V5K1 |
| dnMb7 | Inpaint CD+EH; MPNN | 42% | 48% | UPI001C20C4EB |
| dnMb8 | Inpaint EH; MPNN | 45% | 47% | A0A8C9A9W3 |
| dnMb9 | Inpaint EH; MPNN | 46% | 53% | UPI001C20C4EB |
| dnMb10 | Inpaint EH; MPNN | 46% | 51% | UPI00148EC9BB |
| dnMb11 | Inpaint EH; MPNN | 49% | 55% | P02185 |
| dnMb12 | Inpaint EH; MPNN | 51% | 54% | A0A4W2F1N8 |
| dnMb13 | Inpaint EH; MPNN | 44% | 49% | UPI001CA46E1B |
| dnMb14 | Inpaint EH; MPNN | 46% | 51% | A0A8C9A9W3 |
| dnMb15 | Inpaint EH; MPNN | 46% | 50% | UPI000011026E |
| dnMb16 | Inpaint EH; MPNN | 39% | 45% | P02169 |
| dnMb17 | Inpaint EH; MPNN | 42% | 45% | F6PMG4 |
| dnMb18 | Inpaint CD+EH; MPNN | 42% | 43% | R9RZ90 |
| dnMb19 | Inpaint CD+EH; MPNN | 39% | 41% | UPI0003C8C8C2 |
| dnMb20 | Inpaint CD+EH; MPNN | 45% | 48% | P02182 |
| dnMb21 | Inpaint CD+EH; MPNN | 39% | 44% | P02182 |

**Table S2.** Mass spectrometry data for myoglobin variants.

| Variant | Expected mass | Observed mass |
|---|---|---|
| dnMb2 (Met missing) | 19186 | 19054 |
| dnMb3 (Met missing) | 18981 | 18850 |
| dnMb4 (Met missing) | 19054 | 18923 |
| dnMb5 (Met missing) | 18441 | 18310 |
| dnMb6 (Met missing) | 19604 | 19473 |
| dnMb7 (Met missing) | 18728 | 18597 |
| dnMb8 (Met missing) | 19604 | 19472 |
| dnMb9 (Met missing) | 19579 | 19448 |
| dnMb10 (Met missing) | 18690 | 18559 |
| dnMb11 (Met missing) | 19464 | 19333 |
| dnMb12 (Met missing) | 19536 | 19405 |
| dnMb13 (Met missing) | 19178 | 19047 |
| dnMb14 (Met missing) | 19675 | 19544 |
| dnMb15 (Met partially missing) | 19598 | 19467,19598 |
| dnMb16 (Met partially missing) | 19441 | 19310,19441 |
| dnMb17 (Met missing) | 20247 | 20115 |
| dnMb18 (Met partially missing) | 19427 | 19296,19427 |
| dnMb19 (Met missing) | 19452 | 19321 |
| dnMb20 (Met partially missing) | 18814 | 18683,18814 |
| dnMb21 (Met missing) | 19133 | 19002 |
| Mb 3RGK (Met missing) | 18751 | 18620 |

**Table S3.** The extinction coefficients of the Soret band and $R_z$ values ($A_{Soret}$ / $A_{280}$) of myoglobin variants.

| Variant | Extinction coefficient ($mM^{-1} \cdot cm^{-1}$) | $R_z$ |
|---|---|---|
| dnMb2 | 128 ± 3 | 5.7 |
| dnMb3 | 166 ± 15 | 4.0 |
| dnMb4 | 127 ± 1 | 4.6 |
| dnMb5 | 154 ± 8 | 4.9 |
| dnMb6 | 181 ± 14 | 5.4 |
| dnMb7 | 159 ± 6 | 5.8 |
| dnMb8 | 186 ± 2 | 6.8 |
| dnMb9 | 182 ± 5 | 5.5 |
| dnMb10 | 157 ± 2 | 7.4 |
| dnMb11 | 177 ± 8 | 4.1 |
| dnMb12 | 123 ± 3 | 4.9 |
| dnMb13 | 154 ± 2 | 5.2 |
| dnMb14 | 174 ± 1 | 4.8 |
| dnMb15 | 156 ± 1 | 4.9 |
| dnMb16 | 171 ± 4 | 4.5 |
| dnMb17 | 170 ± 1 | 5.4 |
| dnMb18 | 175 ± 15 | 3.1 |
| dnMb19 | 171 ± 3 | 5.1 |
| dnMb20 | 150 ± 4 | 3.7 |
| dnMb21 | 153 ± 1 | 4.6 |

#### 2. Crystallographic data

Protein sample for crystallography was prepared following the general procedure for myoglobin production. The holoprotein was purified using Ni-affinity and size exclusion chromatography. The C-terminal hexahistidine tag was left intact. The holo dnMb19 was crystallized at 17 mg mL−1 in a buffer containing 25 mM Tris-HCl, 300 mM NaCl, pH 8.2.

Crystallization experiment for the designed protein was conducted using the sitting drop vapor diffusion method. Crystallization trials were set up in 200 nL drops using the 96-well plate format at 20°C. Crystallization plates were set up using a Mosquito LCP from SPT Labtech, then imaged using UVEX microscopes from JAN Scientific. Diffraction quality crystals formed in 0.1 M Bis-Tris pH 6.5, 28% w/v Polyethylene glycol monomethyl ether 2,000 (Index crystallization screen, Hampton Research, well D11).

Diffraction data were collected at ALS-ENABLE beamline 8.2.2. X-ray intensities and data reduction were evaluated and integrated using XDS[1] and merged/scaled using Pointless/Aimless in the CCP4 program suite[2]. Structure determination and refinement starting phases were obtained by molecular replacement using Phaser[3] using the designed model structure. Following molecular replacement, the models were improved using phenix.autobuild[4]. Structures were refined in Phenix[4]. Model building was performed using COOT[5]. The final model was evaluated using MolProbity[6]. Data collection and refinement statistics are recorded in the Supplementary Table 4. Data deposition, atomic coordinates, and structure factors reported for the protein in this paper have been deposited in the Protein Data Bank (PDB), http://www.rcsb.org/ with accession code 8U5A.

**Table S4.** Crystallographic statistics for dnMb19.

| | **dnMb19** |
|---|---|
| PDB accession number | 8U5A |
| Wavelength (Å) | 1.0 |
| Resolution range | 42.64 - 2.0 (2.05 - 2.0) |
| Space group | P 1 $2_1$ 1 |
| Unit cell dimensions<br>a, b, c, (Å)<br>α, β, γ (°) | <br>31.589 41.669 128.439<br>90 95.13 90 |
| Unique reflections | 22200 (1513) |
| Multiplicity | 4.3 (4.1) |
| Completeness (%) | 97.30 (95.52) |
| Mean I/sigma(I) | 10.95 (1.58) |
| Wilson B-factor | 36 |
| R-merge | 0.07531 (0.9919) |
| R-pim | 0.04041 (0.5512) |
| CC1/2 | 0.997 (0.773) |
| Reflections used in refinement | 22200 (1513) |
| R-work | 0.2301 (0.3312) |
| R-free | 0.2581 (0.3741) |
| Number of non-hydrogen atoms | 2612 |
| macromolecules | 2440 |
| ligands | 87 |
| solvent | 85 |
| Protein residues | 298 |
| RMS(bonds) | 0.002 |
| RMS(angles) | 0.39 |
| Ramachandran favored (%) | 99.32 |
| Ramachandran allowed (%) | 0.68 |
| Ramachandran outliers (%) | 0 |
| Average B-factor | 44 |
| macromolecules | 44.03 |
| ligands | 40.92 |
| solvent | 46.5 |

### 3. Sequence information

#### Alignment of TEV hit sequences

```
TEVd            GESLFKGPRDYNPISSTICHLTNESDGHTTSLYGIGFGPFIITNKHLFRRNNGTLLVQSL    60
hyperTEV89      AESAAPGPRDYNPISSTIVRLTNTSDGHSISLFGIGFGPLIITNAHLFRRNNGTLTITSL    60
hyperTEV56      MESAAPGPRDYNPISDTIVKLTNTSDGYSISLYGIGFGPLIITNAHLFRRNNGTLTVTSK    60
hyperTEV60      AESAAPGPRDYNPISDTIVLLTNTSDGYSISLYGIGFGPLIITNAHLFRRNNGTLTITSK    60
                 **   *********.**  *** ***:: **:******:**** ********** : * 

TEVd            HGVFKVKNTTTLQQHLIDGRDMIIIRMPKDFPPFPQKLKFREPQREERICLVTTNFQTKS    120
hyperTEV89      HGTFTISNTTTLKLHLIEGRDLVIIKMPKDFPPFPTTLEFREPVVGEDIVLVTRNFQDKD    120
hyperTEV56      HGTFTIENTTTLQLHLIEGRDLVIIKMPKDFPPFPTDLVFREPVEGEKITLVTRNFQTKE    120
hyperTEV60      HGTFTISNTTTLKLHLIEGRDLVLIEMPKDFPPFPTNLVFREPVVGEEIVLVTRNFQTKT    120
                **.*.:.*****: ***:***:::*.*********  * ****    * * *** *** * 

TEVd            MSSMVSDTSCTFPSSDGIFWKHWIQTKDGQCGSPLVSTRDGFIVGIHSASNFTNTNNYFT    180
hyperTEV89      PTSEVSDTSTTEPSSDGVFWKHWIPTKDGQCGSPMVSVSDGSIVGIHSASNFTNTNNYFT    180
hyperTEV56      PTSEVSDVSSTYPSSDGVFWKHWIPTKDGQCGSPMVSVEDGSIVGIHSASNFTNTNNYFT    180
hyperTEV60      PTSEVSDVSTTYPSSDGVFWKHWIPTKDGQCGSPMVSVTDGSIVGIHSASNFTNTNNYFT    180
                 :* ***.* * *****:****** *********:**. ** ******************

TEVd            SVPKNFMELLTNQEAQQWVSGWRLNADSVLWGGHKVFMDKP   221
hyperTEV89      AVPPNFMDLLTDPSLQKWISGWSLNADSVDWGGHKVFMDKP   221
hyperTEV56      AVPPDFMDLLTNDSLQKWISGWSLNSDSVEWGGHKVFMDKP   221
hyperTEV60      AVPPDFMRLLTDPSLQKWVSGWSLNSDSVEWGGHKVFMDKP   221
                :** :** ***: . *:*:*** **:*** ***********
```

#### Alignment of myoglobin sequences

```
3RGK        -GLSDGEWQLVLNVWGKVEADIPGHGQEVLIRLFKGHPETLEKFDRFKHLKSEDEMKA-S 58
dnMb02      -GLTEEEQKLVWEIFERFEEDLEGFGLDVLIRAFTEHPETLKKFPRFADLKSEAELRA-S 58
dnMb03      -GLTAEEQKLVRDIWAEVEKDREGFGLEVLLLTFTEHPETLKKFPRFAHLKSAEELRA-S 58
dnMb04      -GLTAEEQALVRAIWAKVREDLEGFGLAVLLKTFTEHPETLKKFPRFKDLKSEEEILA-S 58
dnMb05      -GLSDEEQALVLSIFEKVKEDLAGFGLDVLLLAFTKNPATLEKFPRFADLKSEAELLA-S 58
dnMb20      --LSEEEWKIVLEIFALVREDLAGVGAAVLERTFATHPETLKKFPRFLAAAEAGVLD--R 56
dnMb16      ---AEEEKEKVLSIFKLVEKDKKTIGSEVLIITFTKNPETKKKFPRFKDLKTVEELKA-S 56
dnMb19      ---SEEKAALVLALFDRVEADREEIGAAVLRRTFEEHPETLKKFPRFLELYKKGSPEL-D 56
dnMb18      ---DEEKKKLVLEAFELVEKDIEGIGAEVLKLTFEKHPETLEKFPRLKELHAAGSPEL-E 56
dnMb15      ---EEEEKQIVLELFAKVEEDLEGIGLEVLILTFTKHPETRKKFPRFAHLTTEAQLQA-S 56
dnMb17      IKLSEEEKKLVLEIFKLFEENLEEFGKEVLITTFTKHPETKKKFPRFAHLKTEEEFLA-S 59
dnMb13      ----DERNKLVLSAFALVREDLEEIGAEVLILTFTENPETLKKFPRFAHLKTEEELKK-S 55
dnMb14      ---EKEKNELVLKAFELIEKDLEGFGSEVLILTFTKHPETLKKFPRFKHLKTEEEFKA-S 56
dnMb21      -SLTPEELAIVKALFARVREDLEGVGAEVLRLTFEKHPETLKKFPRFLELKKAGSPEL-E 58
dnMb07      -KLTPEEKAIVLRIFALVREDRAGIGAAILRRTFEAHPETLEKFPRLRALRAAGREAELE 59
dnMb06      -KLSEEEKEIVLKIFELVEKDVEEIGLRVLELTFEKHPETLEKFPRLRELLAAGRLEELE 59
dnMb10      ---DAEKQALVASIFAKFEADLEGFGKAVLIKTFTKHPETRKKFPRFKHLKSVEELEK-S 56
dnMb11      -KLSEEEKEIVLKIFALVEKDLEGFGKEVLIKTFLKYPETLKKFPRFKHLKTEEELKA-S 58
dnMb12      LNLSPEDKAKVLEIFALVEEDLEGFGREVLILTFTKHPETLKKFPRFAHLKTEEELRA-S 59
dnMb08      IKISEEEFEIVLEIFELVKKDLAGIGKEVLILTFTKHPETLKKFPRFAHLKTVEELEA-S 59
dnMb09      SKLTEEEWKTVFKIFALVEKDLEGFGLAVLIRTFTRYPETLKKFPRFAHLKTVEELRA-S 59
                      *   :  .. :    *  :*   *   * * :** *:

3RGK        EDLKKHGATVLTALGGI---LKKKGHHEAEIKPLAQSHATKHKIPVKYLEFISEAIIQVL 115
dnMb02      PRLREHGVTVLKALIKI---FKKGEDFAEEVKPLAESHSKVHKIPVSDLEVIAAAILATA 115
dnMb03      PEAKAHGVTVLDALSKI---LKKGSNFEEEIKPLAESHYKEHKIPIEDLKVIADAIIAVL 115
dnMb04      EKAKKHGVTVLTALFAI---FDKGENFEEEIKPLAESHYKEHKIPISDLKVIADAIVAVL 115
dnMb05      EKAKEHGITVLTALFAI---FEKGDDFDAEVEPLATSHTREHKIPTSDLEVIAAAILETA 115
dnMb20      ALLAAHGETVLTALIEIAES----KLDPELIKKLAESHVKEHKIPIEYLRAIADSLIAVL 112
dnMb16      EKVKDHGVTVLDALIEWARLHVEGKDYDSLVKKLAESHKKEHKIPIEDLKSIADALIEVL 116
dnMb19      ALLKEHGKTVLDALIEIARLRYSGEDYRSLIKELAKSHKEEHKIPIEDLRHIAEALLAVL 116
dnMb18      ELLKEHGATVLKALIEIARLKISGGDYLSLVKELAKSHKEEHKIPIEDLKKIAEALLEVL 116
dnMb15      PELKQHGVTVLKALITIAKLYYEGKDYESLIKELAKSHKEEHKIPIEYLEYISESILEVL 116
dnMb17      PELAKHGVTVLNALIEIAKLYLEGKDYRSLIKKLAKSHKLEHKIPIEDLKYIADAIIEVA 119
dnMb13      PLLKEHGVTVLNALIEIAELKYSGGDYESLVKELAKSHKEKHKIPIEDLKAIAEAILKVL 115
dnMb14      EELKEHGVTVLKALIEIAKLKVSGEDYDSLIKELAKSHKTKHKIPIEYLKYIADAILEVA 116
dnMb21      AELRAHGVTVLTALIELADNY---EGNNETLEKLAESHTKVHKIPVSDLKNIAAAIIEVL 115
dnMb07      ALLREHGVTVLDALIEI---V--ENDDEELLKKLAESHKTTHKIPIEHLEHIAAALLEVL 114
dnMb06      AYLREHGVTVLKALIEA---I--KNEDEELLEKLAKSHKEEHKIPIEYLKYIADSIIEVL 114
dnMb10      EELKEHGVTVLTALREIS--L--GENQDKKIKDLATSHKEKHKIPIEDLEVIAAAILEVA 112
dnMb11      EELKEHGVTVLKALIEI---F--KNEDEEKLKELAKSHKEEHKIPIEDLEKIAEAIIEVL 113
dnMb12      EELKEHGVTVLKALRAI---L--EKGDEELLKKLAESHTKEHKIPVSDLEVIAESIIEVA 114
dnMb08      PLLAEHGVTVLKALIKIVEEL--KKGDTSLIKELAKSHKTEHKIDIKDLKYIAESIIEVL 117
dnMb09      PLLREHGVTVLKALTKIAEEL--KKGKTGTLKKLAESHSKVHKIPISDLERIAEAIIEVL 117
                 ** *** **                :. ** **   ***  . *. *: ::: .

3RGK        QSKHPGDFGADAQGAMNKALELFRKDMASNYKEL-    149
dnMb02      KERFPEFFNEKAQAALTKALQQFIDAIAAEYKKL-    149
dnMb03      KKRFPTAFNSAAQAAVTKALQQFIDALEKEFKKL-    149
dnMb04      KEAFPEAFDAKAQAAFTKALEQFIKAFEEEYKKL-    149
dnMb05      KERFPTEFDEEAQAALEKALAQFIAAYAAQAAKL-    149
dnMb20      KERYPERFGEKAQEAVKKFLDLFIEKFEEEAEKEK    147
dnMb16      KKFYPEEFGEEAQAAVQKLLNYFIEKLKQYYE---    148
dnMb19      AERFPDEFGPEARAALTDFLDWFIAEIEEEYKK--    149
dnMb18      KEKYPEEFGEETQEALKEFLDWFIEELEKEFKE--    149
dnMb15      KKRFPEFFGEKAQAAVRKFLDFFISKLKEYYE---    148
dnMb17      KKFFPEKFGEKAQEALKKFLNYFIEELEKEYEKL-    153
dnMb13      KKRYPEEFGEKTQAALKEFLDEFIELTEKYYK---    147
dnMb14      KKRFPKEFDEKTYAALKEFLDYFIEKIEKYYK---    148
dnMb21      KERFPEEFGEEAQAAFTKFLDKFIKDIAELQKKFE    150
dnMb07      AEKYPEEFGPEARAAVTKALELFIKKLAEFYE---    146
dnMb06      EEKFPKEFNEKAREALKKALEYFIEELEKYYK---    146
dnMb10      KERFPEEFDEAAQAALQEFLDDFISKLKEYFE---    144
dnMb11      KEKYPEEFDEEAEEAVKKFLKLFIEKLKEYRE---    145
dnMb12      KKRFPEEFGEEAQAALKKFLEEFIEKWKEYQEEFK    149
dnMb08      KKRFPEEFDEKAKEAVEKVLNLFIEKIEEFYKK--    150
dnMb09      EERFPEEFDEKAKEAVKKFLDLFIEKHAEFVKK--    150
             . .*  *.  :  *. . *  *
```

#### Myoglobin sequences

##### dnMb2

ATGTCAGGAGGCCTGACCGAAGAAGAACAGAAACTGGTGTGGGAAATTTTTGAACGCTTTGAAGAGGATCTGGA
AGGCTTTGGCCTGGATGTGCTGATTCGCGCGTTTACCGAACATCCAGAAACCCTGAAAAAATTTCCGCGCTTTGC
GGATCTGAAAAGCGAAGCGGAATTACGTGCGAGCCCGCGCCTGCGCGAACATGGCGTGACCGTGCTGAAAGCG
CTGATTAAAATCTTTAAAAAAGGCGAAGATTTTGCCGAAGAAGTGAAACCGCTGGCGGAAAGCCATAGCAAAGTG
CATAAAATTCCGGTGAGCGATCTCGAAGTGATTGCGGCGGCGATTCTGGCGACCGCGAAAGAACGCTTTCCGGA
ATTTTTTAATGAAAAAGCGCAGGCGGCGCTGACCAAAGCACTGCAGCAGTTTATTGATGCGATCGCCGCGGAATA
TAAAAAACTGGGCGGCGGTTCCGGCAGCCATCATTGGGGCAGCACCCACCACCACCACCACCAC

MSGGLTEEEQKLVWEIFERFEEDLEGFGLDVLIRAFTEHPETLKKFPRFADLKSEAELRASPRLREHGVTVLKALIKIFK
KGEDFAEEVKPLAESHSKVHKIPVSDLEVIAAAILATAKERFPEFFNEKAQAALTKALQQFIDAIAAEYKKLGGGSGSHH
WGSTHHHHHH

##### dnMb3

ATGTCAGGAGGCCTGACCGCGGAAGAACAGAAACTGGTGCGCGATATTTGGGCGGAAGTGGAAAAAGATCGCG
AAGGCTTTGGCCTGGAAGTGCTGTTGCTGACCTTTACCGAACATCCAGAAACCTTAAAAAAATTTCCACGTTTTGC
GCATCTGAAAAGCGCCGAGGAACTGCGCGCGAGCCCGGAAGCGAAAGCGCATGGCGTGACCGTGCTGGATGC
GCTGAGCAAAATTTTGAAGAAAGGCAGCAATTTCGAAGAAGAAATTAAACCGCTGGCGGAAAGCCATTATAAAGAA
CATAAAATTCCGATTGAAGATTTGAAAGTGATTGCGGATGCGATTATTGCGGTGCTGAAAAAACGCTTTCCGACCG
CGTTTAATAGCGCGGCGCAGGCGGCGGTGACCAAAGCGCTGCAGCAGTTTATTGATGCACTGGAAAAGGAATTT
AAGAAACTGGGCGGCGGTTCCGGCAGCCATCATTGGGGCAGCACCCACCACCACCACCACCAC

MSGGLTAEEQKLVRDIWAEVEKDREGFGLEVLLLTFTEHPETLKKFPRFAHLKSAEELRASPEAKAHGVTVLDALSKIL
KKGSNFEEEIKPLAESHYKEHKIPIEDLKVIADAIIAVLKKRFPTAFNSAAQAAVTKALQQFIDALEKEFKKLGGGSGSHH
WGSTHHHHHH

##### dnMb4

ATGTCAGGAGGTTTAACCGCGGAAGAACAAGCGCTGGTGCGCGCGATTTGGGCGAAAGTGCGCGAAGATCTGG
AAGGCTTTGGCCTGGCGGTGCTGCTGAAAACCTTTACCGAACATCCGGAAACCCTGAAAAAATTTCCGCGCTTTA
AAGACTTGAAAAGCGAAGAAGAAATTCTGGCGAGCGAAAAGGCGAAAAAACATGGCGTGACCGTGCTGACTGCG
CTGTTTGCGATTTTTGATAAAGGTGAGAATTTTGAAGAGGAAATTAAACCGCTGGCGGAAAGCCATTATAAAGAAC
ATAAAATTCCGATTAGCGATCTGAAAGTGATTGCGGATGCGATTGTGGCCGTGTTGAAAGAAGCGTTTCCGGAAG
CATTTGATGCGAAAGCGCAGGCGGCGTTTACCAAAGCACTGGAACAGTTTATTAAAGCGTTCGAGGAAGAATATAA
AAAACTGGGCGGCGGTTCCGGCAGCCATCATTGGGGCAGCACCCACCACCACCACCACCAC

MSGGLTAEEQALVRAIWAKVREDLEGFGLAVLLKTFTEHPETLKKFPRFKDLKSEEEILASEKAKKHGVTVLTALFAIFDK
GENFEEEIKPLAESHYKEHKIPISDLKVIADAIVAVLKEAFPEAFDAKAQAAFTKALEQFIKAFEEEYKKLGGGSGSHHW
GSTHHHHHH

##### dnMb5

ATGTCAGGAGGCCTGAGCGATGAAGAACAGGCGCTGGTGCTGAGCATTTTTGAAAAAGTGAAAGAAGATCTGGC
GGGCTTTGGCCTGGATGTGCTGTTGCTGGCGTTTACCAAAAATCCGGCGACCCTGGAAAAATTTCCGCGCTTTG
CGGATCTGAAAAGCGAAGCGGAACTGTTGGCGAGCGAAAAGGCCAAAGAACATGGCATTACCGTGCTGACCGC
GCTGTTTGCGATTTTCGAGAAAGGCGATGATTTTGATGCGGAAGTTGAACCGCTGGCGACCAGCCATACCCGCG
AACATAAAATTCCGACGAGCGATCTGGAAGTGATTGCGGCGGCGATTCTGGAAACCGCGAAGGAACGCTTTCCA
ACCGAATTTGATGAAGAAGCGCAGGCGGCCTTAGAAAAAGCGTTGGCGCAGTTTATTGCAGCGTATGCGGCGCA
AGCCGCGAAACTGGGCGGCGGTTCCGGCAGCCATCATTGGGGCAGCACCCACCACCACCACCACCAC

MSGGLSDEEQALVLSIFEKVKEDLAGFGLDVLLLAFTKNPATLEKFPRFADLKSEAELLASEKAKEHGITVLTALFAIFEK
GDDFDAEVEPLATSHTREHKIPTSDLEVIAAAILETAKERFPTEFDEEAQAALEKALAQFIAAYAAQAAKLGGGSGSHH
WGSTHHHHHH

**dnMb6**

ATGTCAGGAAAACTGAGCGAAGAAGAAAAAGAAATTGTGCTGAAAATTTTTGAACTGGTGGAAAAGGATGTGGAA
GAAATTGGCCTGCGCGTGCTGGAACTGACCTTTGAAAAACATCCAGAAACCCTGGAGAAATTTCCACGCTTACGC
GAATTATTAGCGGCGGGCCGCCTGGAGGAACTGGAAGCGTATCTGCGCGAACATGGCGTGACCGTGTTAAAAGC
GCTGATTGAAGCGATTAAAAATGAAGATGAAGAACTGTTGGAAAAACTGGCGAAAAGCCATAAAGAGGAACATAAA
ATTCCGATTGAATATCTGAAATATATTGCGGATAGCATTATTGAAGTGTTAGAAGAGAAGTTTCCGAAAGAATTTAAT
GAAAAGGCGCGCGAAGCGTTGAAGAAAGCACTGGAATATTTTATTGAGGAGCTGGAGAAATATTATAAAGGCGGC
GGTTCCGGCAGCCATCATTGGGGCAGCACCCACCACCACCACCACCAC

MSGKLSEEEKEIVLKIFELVEKDVEEIGLRVLELTFEKHPETLEKFPRLRELLAAGRLEELEAYLREHGVTVLKALIEAIKN
EDEELLEKLAKSHKEEHKIPIEYLKYIADSIIEVLEEKFPKEFNEKAREALKKALEYFIEELEKYYKGGGSGSHHWGSTHH
HHHH

**dnMb7**

ATGTCAGGAAAATTAACCCCAGAAGAAAAAGCGATTGTTTTACGTATTTTTGCGTTAGTTCGTGAAGATCGTGCGG
GTATTGGTGCGGCGATTTTGCGTCGTACCTTTGAAGCGCATCCAGAAACCTTAGAAAAATTTCCACGTTTACGTGC
GTTACGCGCCGCGGGCCGCGAAGCGGAACTGGAAGCGCTGTTGCGTGAACATGGCGTGACCGTGCTGGATGC
GCTGATTGAAATTGTGGAAAATGATGATGAAGAACTGCTGAAAAAACTGGCGGAAAGCCATAAAACCACCCACAA
AATTCCAATTGAACATTTAGAACATATTGCGGCGGCGCTGCTGGAAGTGCTGGCCGAGAAATATCCGGAAGAATTT
GGTCCGGAGGCGCGCGCAGCGGTGACCAAAGCCTTGGAACTGTTTATTAAAAAGCTCGCGGAATTTTATGAAGG
CGGCGGTTCCGGCAGCCATCATTGGGGCAGCACCCACCACCACCACCACCAC

MSGKLTPEEKAIVLRIFALVREDRAGIGAAILRRTFEAHPETLEKFPRLRALRAAGREAELEALLREHGVTVLDALIEIVEN
DDEELLKKLAESHKTTHKIPIEHLEHIAAALLEVLAEKYPEEFGPEARAAVTKALELFIKKLAEFYEGGGSGSHHWGSTH
HHHHH

**dnMb8**

ATGTCAGGAATTAAAATTAGCGAAGAAGAATTTGAAATTGTGCTGGAAATTTTTGAACTGGTGAAAAAAGATCTGGC
GGGCATTGGCAAAGAAGTGCTGATTCTGACCTTTACCAAACATCCAGAAACCCTGAAGAAATTTCCACGTTTTGC
GCATCTGAAAACCGTGGAAGAACTGGAAGCGAGCCCGCTGCTGGCGGAACATGGCGTGACCGTGCTGAAAGCG
CTGATTAAGATCGTGGAGGAACTGAAGAAAGGCGATACCAGCCTGATCAAAGAACTGGCGAAAAGCCATAAAACC
GAACATAAGATTGATATTAAGGATTTGAAATATATTGCGGAAAGCATTATTGAAGTTTTAAAAAAACGCTTTCCGGAA
GAGTTCGATGAAAAAGCGAAAGAAGCGGTGGAAAAAGTGTTGAATCTGTTTATCGAGAAAATCGAAGAATTTTATA
AAAAGGGCGGCGGTTCCGGCAGCCATCATTGGGGCAGCACCCACCACCACCACCACCAC

MSGIKISEEEFEIVLEIFELVKKDLAGIGKEVLILTFTKHPETLKKFPRFAHLKTVEELEASPLLAEHGVTVLKALIKIVEELK
KGDTSLIKELAKSHKTEHKIDIKDLKYIAESIIEVLKKRFPEEFDEKAKEAVEKVLNLFIEKIEEFYKKGGGSGSHHWGSTH
HHHHH

**dnMb9**

ATGTCAGGAAGCAAACTGACCGAAGAAGAATGGAAAACCGTGTTTAAAATTTTTGCGCTGGTGGAAAAAGATCTG
GAAGGCTTTGGCCTGGCGGTGCTGATTCGCACCTTTACCCGTTATCCAGAAACCTTGAAAAAATTTCCACGTTTC
GCGCATCTGAAGACCGTGGAAGAATTGCGTGCGAGCCCGCTGCTGCGCGAACATGGCGTGACCGTGCTGAAAG
CGCTGACCAAAATTGCGGAAGAACTGAAGAAAGGCAAAACCGGCACCCTCAAAAAACTGGCGGAAAGCCATAGC
AAAGTGCATAAAATTCCGATTAGCGATTTAGAACGCATTGCCGAAGCGATTATTGAAGTGCTGGAAGAACGCTTTC
CGGAAGAGTTTGATGAAAAAGCGAAAGAAGCGGTGAAGAAGTTTCTGGATCTGTTTATCGAAAAACATGCGGAAT
TTGTGAAAAAAGGCGGCGGTTCCGGCAGCCATCATTGGGGCAGCACCCACCACCACCACCACCAC

MSGSKLTEEEWKTVFKIFALVEKDLEGFGLAVLIRTFTRYPETLKKFPRFAHLKTVEELRASPLLREHGVTVLKALTKIAE
ELKKGKTGTLKKLAESHSKVHKIPISDLERIAEAIIEVLEERFPEEFDEKAKEAVKKFLDLFIEKHAEFVKKGGGSGSHHW
GSTHHHHHH

**dnMb10**

ATGTCAGGAGATGCGGAAAAACAGGCGCTGGTGGCGAGCATTTTTGCGAAATTTGAAGCGGATCTGGAAGGCTT
TGGCAAAGCGGTGCTGATTAAAACCTTTACCAAACATCCGGAAACCCGCAAAAAATTTCCGCGCTTTAAACATCTG
AAAAGCGTGGAAGAACTGGAAAAAAGCGAAGAACTGAAAGAACATGGCGTGACCGTGCTGACCGCGCTGCGCG
AGATTAGCCTGGGCGAAAATCAGGATAAAAAGATTAAAGATCTGGCGACCAGCCATAAAGAAAAGCATAAAATTCC
GATTGAAGATTTGGAAGTGATTGCGGCGGCGATTTTAGAAGTGGCGAAGGAACGCTTTCCGGAAGAATTTGATGA
GGCGGCGCAGGCAGCGCTGCAGGAATTTCTGGATGATTTTATTAGCAAATTAAAAGAATATTTTGAAGGCGGCGG
TTCCGGCAGCCATCATTGGGGCAGCACCCACCACCACCACCACCAC

MSGDAEKQALVASIFAKFEADLEGFGKAVLIKTFTKHPETRKKFPRFKHLKSVEELEKSEELKEHGVTVLTALREISLGE
NQDKKIKDLATSHKEKHKIPIEDLEVIAAAILEVAKERFPEEFDEAAQAALQEFLDDFISKLKEYFEGGGSGSHHWGSTH
HHHHH

**dnMb11**

ATGTCAGGAAAACTGAGCGAAGAAGAAAAAGAAATTGTGCTGAAAATTTTTGCGCTGGTGGAAAAGGATCTGGAA
GGCTTTGGCAAAGAAGTGCTGATTAAAACCTTTCTGAAATATCCGGAAACCCTGAAAAAATTTCCGCGCTTTAAAC
ATCTGAAAACCGAGGAAGAACTGAAAGCGTCGGAAGAGTTGAAAGAACATGGCGTGACCGTGTTAAAAGCGCTG
ATTGAAATCTTTAAAAATGAAGATGAAGAAAAACTGAAGGAGCTGGCGAAAAGCCATAAAGAAGAGCATAAAATTC
CGATTGAAGATTTAGAGAAAATTGCGGAAGCGATTATTGAAGTACTGAAAGAGAAATACCCGGAAGAATTTGATGA
AGAAGCGGAGGAGGCGGTGAAAAAGTTTTTAAAACTGTTTATCGAGAAGCTCAAAGAATATCGCGAAGGCGGCG
GTTCCGGCAGCCATCATTGGGGCAGCACCCACCACCACCACCACCAC

MSGKLSEEEKEIVLKIFALVEKDLEGFGKEVLIKTFLKYPETLKKFPRFKHLKTEEELKASEELKEHGVTVLKALIEIFKNE
DEEKLKELAKSHKEEHKIPIEDLEKIAEAIIEVLKEKYPEEFDEEAEEAVKKFLKLFIEKLKEYREGGGSGSHHWGSTHH
HHHH

**dnMb12**

ATGTCAGGACTGAATCTGAGCCCGGAAGATAAAGCGAAAGTGCTGGAAATTTTTGCGCTGGTGGAAGAAGATCTG
GAAGGCTTTGGCCGCGAAGTTCTGATTCTGACCTTTACCAAACATCCGGAAACCCTGAAAAAATTTCCACGCTTT
GCGCATCTGAAAACCGAAGAGGAACTGCGCGCGAGCGAAGAACTGAAAGAACATGGCGTGACCGTGCTGAAAG
CGCTGCGTGCGATTCTGGAAAAAGGCGATGAAGAGCTGTTGAAGAAACTGGCGGAAAGCCATACCAAAGAACAT
AAAATTCCGGTGAGCGATTTGGAAGTGATTGCGGAAAGCATTATTGAAGTGGCGAAAAAACGCTTTCCGGAGGAA
TTTGGTGAAGAAGCGCAGGCGGCGCTGAAGAAGTTTTTAGAAGAATTTATCGAAAAATGGAAAGAATATCAGGAG
GAGTTTAAAGGCGGCGGTTCCGGCAGCCATCATTGGGGCAGCACCCACCACCACCACCACCAC

MSGLNLSPEDKAKVLEIFALVEEDLEGFGREVLILTFTKHPETLKKFPRFAHLKTEEELRASEELKEHGVTVLKALRAILE
KGDEELLKKLAESHTKEHKIPVSDLEVIAESIIEVAKKRFPEEFGEEAQAALKKFLEEFIEKWKEYQEEFKGGGSGSHH
WGSTHHHHHH

**dnMb13**

ATGTCAGGAGATGAACGCAATAAACTGGTGCTGAGCGCGTTTGCGCTGGTGCGCGAAGATCTGGAAGAAATTGG
CGCGGAAGTGCTGATTCTGACCTTTACCGAAAATCCGGAAACCCTGAAAAAATTTCCGCGCTTTGCGCATCTGAA
AACCGAAGAAGAACTGAAAAAAAGCCCGCTGCTGAAAGAACATGGCGTGACCGTGCTGAATGCGCTGATTGAAA
TTGCGGAATTAAAATATAGCGGCGGCGATTATGAAAGCCTGGTGAAAGAGCTGGCGAAAAGCCATAAAGAAAAAC
ATAAAATTCCGATTGAAGATTTGAAAGCGATTGCAGAAGCCATCTTGAAAGTCTTAAAAAAACGCTATCCAGAAGAA

TTTGGTGAGAAGACCCAGGCGGCGTTGAAGGAGTTTCTGGATGAATTTATTGAACTGACCGAGAAATATTATAAAG
GCGGCGGTTCCGGCAGCCATCATTGGGGCAGCACCCACCACCACCACCACCAC

MSGDERNKLVLSAFALVREDLEEIGAEVLILTFTENPETLKKFPRFAHLKTEEELKKSPLLKEHGVTVLNALIEIAELKYSG
GDYESLVKELAKSHKEKHKIPIEDLKAIAEAILKVLKKRYPEEFGEKTQAALKEFLDEFIELTEKYYKGGGSGSHHWGST
HHHHHH

**dnMb14**

ATGTCAGGAGAAAAAGAAAAAAATGAACTGGTGCTGAAAGCGTTTGAACTGATTGAAAAGGATCTGGAAGGCTTT
GGCAGCGAAGTGCTGATTCTGACCTTTACCAAACATCCGGAAACCCTGAAAAAATTTCCGCGCTTTAAACATCTGA
AAACCGAAGAAGAATTTAAAGCGAGCGAAGAACTGAAAGAACATGGCGTGACCGTGTTGAAGGCGCTGATCGAA
ATTGCGAAATTAAAAGTGAGCGGCGAAGATTATGATAGCCTGATTAAAGAGCTGGCGAAAAGCCATAAAACCAAAC
ATAAAATTCCGATTGAATATTTGAAATATATTGCGGATGCGATTTTGGAAGTGGCAAAAAAACGCTTTCCGAAAGAG
TTTGATGAGAAGACCTATGCGGCGTTAAAGGAATTTCTGGATTATTTTATTGAGAAAATTGAGAAATATTATAAAGGC
GGCGGTTCCGGCAGCCATCATTGGGGCAGCACCCACCACCACCACCACCAC

MSGEKEKNELVLKAFELIEKDLEGFGSEVLILTFTKHPETLKKFPRFKHLKTEEEFKASEELKEHGVTVLKALIEIAKLKVS
GEDYDSLIKELAKSHKTKHKIPIEYLKYIADAILEVAKKRFPKEFDEKTYAALKEFLDYFIEKIEKYYKGGGSGSHHWGST
HHHHHH

**dnMb15**

ATGTCAGGAGAAGAAGAAGAAAAACAGATTGTGCTGGAACTGTTTGCGAAAGTGGAAGAGGATCTGGAAGGCAT
TGGCCTGGAAGTGCTGATTCTGACCTTTACCAAACATCCAGAAACCCGTAAAAAATTTCCACGTTTTGCGCATCTG
ACCACCGAAGCGCAGCTGCAGGCGAGCCCGGAACTGAAACAGCATGGCGTGACCGTGCTGAAAGCGCTGATTA
CCATTGCCAAACTGTATTATGAAGGCAAAGATTACGAAAGCCTGATTAAAGAACTGGCGAAAAGCCATAAAGAGGA
ACATAAAATTCCGATTGAATATTTGGAATATATTAGCGAAAGCATTTTGGAGGTTCTGAAAAAACGCTTTCCGGAATT
TTTTGGTGAGAAAGCCCAGGCGGCGGTGCGCAAATTTCTGGATTTTTTTATTAGCAAATTGAAAGAATATTACGAA
GGCGGCGGTTCCGGCAGCCATCATTGGGGCAGCACCCACCACCACCACCACCAC

MSGEEEEKQIVLELFAKVEEDLEGIGLEVLILTFTKHPETRKKFPRFAHLTTEAQLQASPELKQHGVTVLKALITIAKLYYE
GKDYESLIKELAKSHKEEHKIPIEYLEYISESILEVLKKRFPEFFGEKAQAAVRKFLDFFISKLKEYYEGGGSGSHHWGS
THHHHHH

**dnMb16**

ATGTCAGGAGCGGAAGAAGAAAAAGAAAAAGTGCTGAGCATTTTTAAACTGGTGGAAAAGGATAAAAAAACCATT
GGCAGCGAAGTCCTGATTATTACCTTTACCAAAAATCCGGAAACCAAGAAGAAATTTCCGCGCTTTAAAGATCTGA
AAACCGTGGAAGAACTGAAAGCGAGCGAAAAAGTGAAAGATCATGGCGTGACCGTGCTGGATGCGCTGATTGAA
TGGGCGCGCCTGCATGTGGAAGGCAAAGATTATGATAGCCTGGTGAAAAAACTGGCGGAAAGCCATAAGAAGGA
ACATAAAATTCCGATTGAAGATTTGAAAAGCATTGCGGACGCCTTGATCGAAGTTTTAAAGAAATTTTATCCAGAGG
AATTTGGCGAAGAAGCGCAGGCGGCGGTGCAGAAACTGCTGAATTATTTTATTGAGAAGTTGAAACAGTATTATGA
AGGCGGCGGTTCCGGCAGCCATCATTGGGGCAGCACCCACCACCACCACCACCAC

MSGAEEEKEKVLSIFKLVEKDKKTIGSEVLIITFTKNPETKKKFPRFKDLKTVEELKASEKVKDHGVTVLDALIEWARLHV
EGKDYDSLVKKLAESHKKEHKIPIEDLKSIADALIEVLKKFYPEEFGEEAQAAVQKLLNYFIEKLKQYYEGGGSGSHHW
GSTHHHHHH

**dnMb17**

ATGTCAGGAATTAAACTGAGCGAAGAAGAAAAAAAACTGGTGCTGGAAATTTTTAAGCTGTTTGAAGAGAATCTGG
AAGAATTTGGCAAAGAAGTGCTGATTACCACCTTTACCAAACATCCGGAAACCAAAAAAAAATTTCCGCGCTTTGC
GCATCTGAAAACCGAGGAGGAATTTCTGGCGAGCCCGGAACTGGCGAAACATGGCGTGACCGTGCTGAATGCG
CTGATTGAAATTGCGAAACTGTATTTAGAAGGCAAGGATTATCGCAGCCTGATTAAAAAGCTGGCAAAAAGCCATA

AACTGGAACATAAAATTCCGATTGAAGATTTGAAATATATTGCGGATGCGATTATTGAAGTGGCCAAAAAGTTCTTTC
CAGAAAAATTCGGTGAAAAAGCGCAGGAAGCGCTGAAGAAGTTCCTGAATTATTTTATTGAGGAGTTGGAAAAAG
AATATGAAAAACTGGGCGGCGGTTCCGGCAGCCATCATTGGGGCAGCACCCACCACCACCACCACCAC

MSGIKLSEEEKKLVLEIFKLFEENLEEFGKEVLITTFTKHPETKKKFPRFAHLKTEEEFLASPELAKHGVTVLNALIEIAKLY
LEGKDYRSLIKKLAKSHKLEHKIPIEDLKYIADAIIEVAKKFFPEKFGEKAQEALKKFLNYFIEELEKEYEKLGGGSGSHH
WGSTHHHHHH

#### dnMb18

ATGTCAGGAGATGAAGAAAAGAAAAAACTGGTGCTGGAAGCGTTTGAATTGGTGGAAAAAGATATTGAAGGCATT
GGCGCGGAAGTGCTGAAACTGACCTTTGAAAAACATCCGGAAACCCTGGAGAAATTTCCGCGCCTGAAAGAATTA
CATGCGGCGGGCAGCCCGGAACTGGAAGAACTGTTGAAGGAACATGGCGCGACCGTGTTGAAAGCGCTGATTG
AAATTGCGCGCTTAAAAATTAGCGGCGGCGATTATCTGAGCCTGGTGAAAGAGCTGGCGAAAAGCCATAAAGAAG
AGCATAAAATTCCGATTGAAGATCTGAAGAAAATCGCCGAAGCCCTGCTGGAGGTTTTGAAGGAAAAATATCCAGA
AGAATTTGGCGAAGAAACCCAGGAGGCTCTGAAAGAATTTCTGGATTGGTTTATCGAGGAGCTGGAAAAGGAATT
TAAGGAAGGCGGCGGTTCCGGCAGCCATCATTGGGGCAGCACCCACCACCACCACCACCAC

MSGDEEKKKLVLEAFELVEKDIEGIGAEVLKLTFEKHPETLEKFPRLKELHAAGSPELEELLKEHGATVLKALIEIARLKIS
GGDYLSLVKELAKSHKEEHKIPIEDLKKIAEALLEVLKEKYPEEFGEETQEALKEFLDWFIEELEKEFKEGGGSGSHHW
GSTHHHHHH

#### dnMb19

ATGTCAGGAAGCGAAGAAAAAGCGGCGTTGGTGCTGGCGCTGTTTGATCGCGTGGAAGCGGATCGCGAAGAAA
TTGGTGCGGCGGTGCTGCGCCGCACCTTTGAAGAACATCCGGAAACCCTGAAAAAATTTCCGCGCTTTCTGGAA
CTGTATAAAAAAGGTAGCCCAGAACTGGATGCGCTGCTGAAAGAGCATGGCAAAACCGTGCTGGACGCGCTGAT
TGAAATTGCGCGCCTGCGCTATAGCGGCGAAGATTATCGCAGCCTGATTAAAGAGCTGGCGAAAAGCCATAAAGA
AGAACATAAAATTCCAATTGAAGATCTGCGCCATATTGCGGAAGCGTTGTTGGCCGTTTTGGCGGAACGCTTTCC
GGATGAATTTGGTCCAGAAGCGCGCGCGGCACTGACCGATTTTTTGGATTGGTTTATCGCGGAAATTGAGGAGGA
ATATAAGAAAGGCGGCGGTTCCGGCAGCCATCATTGGGGCAGCACCCACCACCACCACCACCAC

MSGSEEKAALVLALFDRVEADREEIGAAVLRRTFEEHPETLKKFPRFLELYKKGSPELDALLKEHGKTVLDALIEIARLR
YSGEDYRSLIKELAKSHKEEHKIPIEDLRHIAEALLAVLAERFPDEFGPEARAALTDFLDWFIAEIEEEYKKGGGSGSHH
WGSTHHHHHH

#### dnMb20

ATGTCAGGACTGAGCGAAGAAGAATGGAAAATTGTGCTGGAAATTTTTGCGTTAGTTCGTGAAGATTTAGCGGGT
GTTGGTGCGGCGGTTTTAGAACGTACCTTTGCGACCCATCCAGAAACCTTAAAAAAATTTCCACGTTTTCTCGCG
GCAGCGGAAGCGGGCGTGTTGGATCGTGCGCTGCTGGCCGCGCATGGCGAAACCGTGCTGACCGCGCTGATT
GAAATTGCGGAAAGCAAACTGGATCCGGAACTGATTAAGAAACTGGCGGAAAGCCATGTGAAAGAACATAAAATT
CCGATTGAATATCTGCGCGCGATTGCCGATAGCCTGATCGCGGTCCTGAAAGAACGCTATCCAGAGCGCTTTGGT
GAAAAAGCGCAGGAGGCGGTTAAAAAGTTTCTGGATCTGTTTATTGAAAAGTTTGAAGAAGAAGCGGAAAAAGAA
AAAGGCGGCGGTTCCGGCAGCCATCATTGGGGCAGCACCCACCACCACCACCACCAC

MSGLSEEEWKIVLEIFALVREDLAGVGAAVLERTFATHPETLKKFPRFLAAAEAGVLDRALLAAHGETVLTALIEIAESKL
DPELIKKLAESHVKEHKIPIEYLRAIADSLIAVLKERYPERFGEKAQEAVKKFLDLFIEKFEEEAEKEKGGGSGSHHWGS
THHHHHH

#### dnMb21

ATGTCAGGAAGCTTAACCCCAGAAGAATTAGCGATTGTGAAAGCGCTGTTTGCGCGCGTGCGCGAAGATCTGGA
AGGCGTGGGCGCGGAAGTGCTGCGCTTGACCTTTGAAAAACATCCAGAAACCTTAAAAAAATTTCCACGTTTTTT
GGAATTGAAGAAAGCGGGCAGCCCGGAACTGGAAGCGGAGCTGCGTGCGCATGGCGTGACCGTGCTGACCGC

GCTGATTGAACTTGCGGATAATTATGAAGGCAATAATGAAACTCTGGAAAAACTGGCCGAAAGCCATACCAAAGTG
CATAAAATTCCGGTGAGCGATCTGAAGAATATTGCGGCGGCGATTATTGAAGTCCTGAAAGAACGCTTTCCGGAA
GAGTTTGGCGAAGAAGCGCAGGCGGCGTTTACCAAATTTTTAGATAAATTTATTAAAGATATTGCGGAGCTGCAGA
AAAAATTTGAAGGCGGCGGTTCCGGCAGCCATCATTGGGGCAGCACCCACCACCACCACCAC

MSGSLTPEELAIVKALFARVREDLEGVGAEVLRLTFEKHPETLKKFPRFLELKKAGSPELEAELRAHGVTVLTALIELAD
NYEGNNETLEKLAESHTKVHKIPVSDLKNIAAAIIEVLKERFPEEFGEEAQAAFTKFLDKFIKDIAELQKKFEGGGSGSH
HWGSTHHHHHH

##### Native Mb, 3RGK

ATGTCAGGAGGCCTGAGCGATGGCGAATGGCAGCTGGTGCTGAACGTGTGGGGCAAAGTGGAAGCGGATATTC
CGGGCCATGGCCAGGAAGTGCTGATTCGCCTGTTTAAAGGTCATCCGGAAACCCTGGAAAAGTTCGACCGCTTT
AAACATCTGAAATCTGAAGATGAGATGAAAGCGAGCGAAGATCTGAAGAAACATGGCGCGACCGTGCTGACCGC
GCTGGGCGGTATTCTGAAGAAGAAAGGCCATCACGAAGCCGAAATTAAACCGCTGGCGCAGAGCCATGCGACCA
AACATAAAATTCCGGTGAAGTACCTGGAATTTATCAGCGAAGCGATTATTCAGGTGCTGCAGAGCAAACATCCGGG
CGATTTTGGCGCGGATGCGCAGGGTGCGATGAACAAAGCGCTGGAACTGTTTCGCAAAGATATGGCGAGCAACT
ATAAAGAACTGGGTTCCGGCAGCCATCATTGGGGCAGCACCCACCACCACCACCAC

MSGGLSDGEWQLVLNVWGKVEADIPGHGQEVLIRLFKGHPETLEKFDRFKHLKSEDEMKASEDLKKHGATVLTALGGI
LKKKGHHEAEIKPLAQSHATKHKIPVKYLEFISEAIIQVLQSKHPGDFGADAQGAMNKALELFRKDMASNYKELGSGSH
HWGSTHHHHHH

##### TEV sequences

N-terminal tag + linker and C-terminal linker are highlighted in green.

##### hyperTEV56

ATGAGCCACCACCACCACCACCACTCAGGAATGGAAAGCGCGGCGCCGGGCCCGCGCGATTATAACCCGATCA
GCGATACCATTGTGAAACTGACCAACACCTCTGATGGCTATAGCATTAGCCTGTATGGCATTGGCTTTGGCCCGCT
GATCATTACCAACGCGCATTTATTTCGCCGCAACAACGGCACCCTGACCGTGACCAGCAAACATGGCACCTTCAC
CATTGAAAACACCACCACCCTGCAGCTGCATCTGATTGAAGGCCGCGATCTGGTGATTATTAAAATGCCGAAAGAT
TTTCCGCCGTTTCCGACCGATCTGGTGTTTCGCGAACCGGTGGAAGGCGAGAAAATTACCCTGGTGACCCGCAA
CTTTCAGACCAAAGAACCGACCAGCGAAGTGAGCGATGTGAGCAGCACCTATCCGAGCAGCGATGGCGTGTTTT
GGAAACATTGGATTCCGACCAAAGATGGTCAGTGCGGCAGCCCGATGGTGAGCGTGGAAGATGGCAGCATTGTG
GGCATTCATAGCGCGAGCAACTTTACCAATACCAACAACTATTTTACCGCGGTGCCGCCGGATTTCATGGATCTGC
TGACCAACGATAGCCTGCAGAAATGGATTAGCGGCTGGAGCCTGAACAGCGATAGCGTTGAATGGGGCGGCCAT
AAAGTGTTTATGGATAAACCGGGTTCC

MSHHHHHHSGMESAAPGPRDYNPISDTIVKLTNTSDGYSISLYGIGFGPLIITNAHLFRRNNGTLTVTSKHGTFTIENTT
TLQLHLIEGRDLVIIKMPKDFPPFPTDLVFREPVEGEKITLVTRNFQTKEPTSEVSDVSSTYPSSDGVFWKHWIPTKDGQ
CGSPMVSVEDGSIVGIHSASNFTNTNNYFTAVPPDFMDLLTNDSLQKWISGWSLNSDSVEWGGHKVFMDKPGS

##### hyperTEV60

ATGAGCCACCACCACCACCACCACTCAGGAGCGGAAAGCGCGGCGCCGGGCCCGCGCGATTATAACCCGATTA
GCGATACCATTGTTCTGCTGACCAATACCAGCGATGGCTATAGCATTAGCCTGTATGGCATTGGCTTTGGCCCGCT
GATTATTACCAACGCGCACCTGTTTCGCCGCAACAACGGCACCCTGACCATTACCAGCAAACATGGCACCTTTAC
CATTAGCAACACCACCACCCTGAAACTGCATCTGATCGAAGGCCGCGATCTGGTGCTGATTGAAATGCCGAAAGA
TTTTCCGCCGTTTCCGACCAACCTGGTGTTTCGTGAACCGGTGGTGGGCGAAGAAATTGTGCTGGTGACCCGCA
ACTTTCAGACCAAAACCCCGACCAGCGAAGTGAGCGATGTGAGCACCACCTATCCGAGCTCCGATGGCGTGTTT
TGGAAACATTGGATTCCGACGAAAGATGGCCAGTGCGGCAGCCCGATGGTGAGCGTGACCGATGGCAGCATTGT
GGGCATTCATAGCGCGAGCAACTTTACCAACACCAACAACTATTTTACCGCGGTGCCGCCGGATTTTATGCGCCT
GCTGACCGATCCGAGCCTGCAGAAATGGGTGAGCGGCTGGAGCCTGAACAGCGATAGCGTGGAATGGGGCGG
CCATAAAGTGTTTATGGATAAACCGGGTTCC

MSHHHHHHSGAESAAPGPRDYNPISDTIVLLTNTSDGYSISLYGIGFGPLIITNAHLFRRNNGTLTITSKHGTFTISNTTTL
KLHLIEGRDLVLIEMPKDFPPFPTNLVFREPVVGEEIVLVTRNFQTKTPTSEVSDVSTTYPSSDGVFWKHWIPTKDGQC
GSPMVSVTDGSIVGIHSASNFTNTNNYFTAVPPDFMRLLTDPSLQKWVSGWSLNSDSVEWGGHKVFMDKPGS

##### hyperTEV89

ATGAGCCACCACCACCACCACCACTCAGGAGCGGAAAGCGCGGCGCCGGGCCCGCGCGATTATAACCCGATTA
GCAGCACCATTGTGCGCCTGACCAACACCAGCGACGGCCATAGCATTAGCCTGTTTGGCATTGGCTTTGGCCCG
CTGATTATTACCAACGCGCATTTATTTCGCCGCAACAACGGCACCCTGACCATTACCAGCCTGCATGGCACCTTTA
CCATTAGCAACACCACCACCCTGAAACTGCATCTGATTGAAGGCCGCGATCTGGTGATTATCAAAATGCCGAAAG
ATTTTCCGCCGTTTCCGACCACCCTGGAATTTCGCGAACCGGTGGTGGGCGAAGATATTGTGCTGGTGACCCGC
AACTTTCAGGATAAAGATCCGACCAGCGAAGTGAGCGATACCAGCACCACCGAACCGAGCAGTGATGGCGTGTT
TTGGAAACATTGGATCCCGACCAAAGATGGCCAGTGCGGCAGCCCGATGGTGAGCGTTAGCGATGGCAGCATTG
TGGGCATTCATAGCGCGAGCAACTTCACCAATACCAACAACTATTTTACCGCGGTGCCGCCGAACTTTATGGATCT
GCTGACCGATCCGAGCCTGCAGAAATGGATTAGCGGCTGGAGCCTGAACGCGGATAGCGTGGATTGGGGCGGC
CATAAAGTGTTCATGGATAAACCGGGTTCC

MSHHHHHHSGAESAAPGPRDYNPISSTIVRLTNTSDGHSISLFGIGFGPLIITNAHLFRRNNGTLTITSLHGTFTISNTTTL
KLHLIEGRDLVIIKMPKDFPPFPTTLEFREPVVGEDIVLVTRNFQDKDPTSEVSDTSTTEPSSDGVFWKHWIPTKDGQC
GSPMVSVSDGSIVGIHSASNFTNTNNYFTAVPPNFMDLLTDPSLQKWISGWSLNADSVDWGGHKVFMDKPGS

##### TEVd (PDB: 1LVM)[7]

ATGAGCCACCACCACCACCACCACTCAGGAGGCGAAAGCCTGTTCAAAGGCCCGCGCGATTATAACCCGATTAG
CAGCACCATTTGCCATCTGACCAACGAAAGCGATGGCCATACCACCAGCCTGTATGGCATTGGCTTTGGCCCGTT
TATCATTACCAACAAACATTTATTTCGCCGCAACAACGGCACCCTGCTGGTGCAGAGCCTGCATGGCGTGTTTAAA
GTGAAGAATACCACGACCCTGCAGCAGCATCTGATTGATGGCCGCGATATGATTATTATTCGCATGCCGAAAGATT
TTCCGCCGTTTCCGCAGAAACTGAAATTTCGCGAACCGCAGCGTGAAGAACGCATTTGTCTGGTGACCACCAAC
TTTCAGACGAAAAGCATGAGCAGCATGGTGAGCGATACCAGCTGCACCTTTCCGAGCAGCGATGGTATCTTTTGG
AAACATTGGATTCAGACCAAAGATGGTCAGTGCGGCAGCCCGCTGGTGAGCACCCGTGATGGCTTTATTGTGGG
CATTCATAGCGCGAGCAACTTTACCAATACCAATAACTATTTTACCAGCGTGCCGAAGAACTTTATGGAACTGCTGA
CCAACCAGGAAGCGCAGCAGTGGGTGAGCGGCTGGCGCCTGAACGCGGATAGCGTGCTGTGGGGCGGCCATA
AAGTGTTTATGGATAAACCGGGTTCC

MSHHHHHHSGGESLFKGPRDYNPISSTICHLTNESDGHTTSLYGIGFGPFIITNKHLFRRNNGTLLVQSLHGVFKVKNT
TTLQQHLIDGRDMIIIRMPKDFPPFPQKLKFREPQREERICLVTTNFQTKSMSSMVSDTSCTFPSSDGIFWKHWIQTKD
GQCGSPLVSTRDGFIVGIHSASNFTNTNNYFTSVPKNFMELLTNQEAQQWVSGWRLNADSVLWGGHKVFMDKPGS

### S219V[7]

ATGAGCCACCACCACCACCACCACTCAGGAGGCGAAAGCCTGTTCAAAGGCCCGCGCGATTATAACCCGATTAG
CAGCACCATTTGCCATCTGACCAATGAAAGCGATGGCCATACCACCAGCCTGTATGGCATTGGCTTTGGCCCGTT
TATTATTACCAACAAACATTTATTTCGCCGCAACAACGGCACCCTGCTGGTGCAGAGCCTGCATGGCGTGTTTAAA
GTTAAGAACACCACCACCCTGCAGCAGCATCTGATTGATGGCCGCGATATGATCATTATTCGCATGCCGAAAGATT
TTCCGCCGTTTCCGCAGAAACTGAAATTTCGCGAACCGCAGCGCGAAGAACGCATTTGCCTGGTGACCACCAAC
TTTCAGACCAAAAGCATGAGCAGCATGGTGAGCGATACCAGCTGCACCTTTCCGAGCAGCGATGGTATCTTTTGG
AAACATTGGATTCAGACGAAAGATGGCCAGTGCGGCAGCCCGCTGGTGAGCACCCGTGATGGCTTTATTGTGGG
CATTCATAGCGCGAGCAACTTTACCAACACCAACAACTATTTTACCAGCGTGCCGAAGAATTTTATGGAACTGCTG
ACCAACCAGGAAGCGCAGCAGTGGGTGAGCGGCTGGCGCCTGAACGCGGATAGCGTGCTGTGGGGCGGCCAT
AAAGTGTTTATGGTGAAACCGGAAGAACCGTTTCAGCCGGTGAAAGAAGCGACCCAGCTGATGAACGAAGGTTC
C

MSHHHHHHSGGESLFKGPRDYNPISSTICHLTNESDGHTTSLYGIGFGPFIITNKHLFRRNNGTLLVQSLHGVFKVKNT
TTLQQHLIDGRDMIIIRMPKDFPPFPQKLKFREPQREERICLVTTNFQTKSMSSMVSDTSCTFPSSDGIFWKHWIQTKD
GQCGSPLVSTRDGFIVGIHSASNFTNTNNYFTSVPKNFMELLTNQEAQQWVSGWRLNADSVLWGGHKVFMVKPEEP
FQPVKEATQLMNEGS

#### TEV1Δ[8]

ATGAGCCACCACCACCACCACCACTCAGGAGGCGAAAGCCTGTTTAAAGGCCCGCGCGATTATAACCCGATTAG
CAGCACCATTTGCCATCTGACCAACGAAAGCGATGGCCATACCACCAGCCTGTATGGCATTGGCTTTGGCCCGTT
TATTATTACCAATAAACATTTATTTCGCCGCAACAACGGCACCCTGCTGGTGCAGAGCCTGCATGGCGTGTTTAAA
GTGAAGAATACCACGACCCTGCAGCAGCATCTGATTGATGGCCGCGATATGATTATTATTCGCATGCCGAAAGATT
TTCCGCCGTTTCCGCAGAAACTGAAATTTCGCGAACCGCAGCGCGAAGAACGCATCTGCCTGGTGACCACCAAC
TTTCAGACCAAAAGCATGAGCAGCATGGTGAGCGATACCAGCTGCACCTTTCCGAGCAGCGACGGCATTTTCTG
GAAACATTGGATTCAGACGAAAGATGGTCAGTGCGGCAACCCGCTGGTGAGCACCCGCGATGGCTTTATTGTGG
GCATTCATAGCGCGAGCAACTTTACCAACACCAACAACTATTTTACCAGCGTGCCGAAGAACTTTATGGAACTGCT
GACCAACCAGGAAGCGCAGCAGTGGGTGAGCGGCTGGCGCCTGAACGCGGATAGCGTGCTGTGGGGCGGCCA
TAAAGTGTTTATGGTGGGTTCC

MSHHHHHHSGGESLFKGPRDYNPISSTICHLTNESDGHTTSLYGIGFGPFIITNKHLFRRNNGTLLVQSLHGVFKVKNT
TTLQQHLIDGRDMIIIRMPKDFPPFPQKLKFREPQREERICLVTTNFQTKSMSSMVSDTSCTFPSSDGIFWKHWIQTKD
GQCGNPLVSTRDGFIVGIHSASNFTNTNNYFTSVPKNFMELLTNQEAQQWVSGWRLNADSVLWGGHKVFMVGS

#### superTEV[9]

ATGAGCCACCACCACCACCACCACTCAGGACCGCGCGATTATAACCCGATTAGCAGCACCATTGTGCATCTGACC
AACGAAAGCGATGGCCATACCACCAGCCTGTATGGCATTGGCTTTGGCCCGTTTATTATTACCAACAAACATTTATT
TCGCCGCAACAACGGCACCCTGCTGGTGCAGAGCCTGCATGGCGTGTTTAAAGTTAAGAACACCACCACCCTGC
AGCAGCATCTGATTGATGGCCGCGATATGATTATTATCCGCATGCCGAAAGATTTTCCGCCGTTTCCGCAGAAACT
GAAATTTCGCGAACCGCAGCGTGAAGAACGTATTGTGCTGGTGACCACCAACTTTCAGACCAAAAGCATGAGCA
GCATGGTGAGCGATACCAGCAGCACCTTTCCGAGCAGCGATGGTATTTTCTGGAAACATTGGATCCAGACCAAAG
ATGGCCAGTGCGGCAGCCCGCTGGTGAGCACCCGTGATGGCTTTATTGTGGGCATTCATAGCGCGAGCAACTTT
ACCAACACCAATAACTATTTTACCAGCGTGCCGAAGAACTTTATGGAACTGCTGACCAATCAGGAAGCGCAGCAG
TGGGTGAGCGGCTGGCGCCTGAACGCGGATAGCGTGCTGTGGGGCGGCCATAAAGTGTTTATGGATAAACCGG
GTTCC

MSHHHHHHSGPRDYNPISSTIVHLTNESDGHTTSLYGIGFGPFIITNKHLFRRNNGTLLVQSLHGVFKVKNTTTLQQHLI
DGRDMIIIRMPKDFPPFPQKLKFREPQREERIVLVTTNFQTKSMSSMVSDTSSTFPSSDGIFWKHWIQTKDGQCGSPL
VSTRDGFIVGIHSASNFTNTNNYFTSVPKNFMELLTNQEAQQWVSGWRLNADSVLWGGHKVFMDKPGS

#### MBP-TEVcs-FKBP-EGFP substrate

TEVcs is highlighted in orange, FKBP is highlighted in green, and EGFP is highlighted in yellow.

MKIEEGKLVIWINGDKGYNGLAEVGKKFEKDTGIKVTVEHPDKLEEKFPQVAATGDGPDIIFWAHDRFGGYAQSGLLAE
ITPDKAFQDKLYPFTWDAVRYNGKLIAYPIAVEALSLIYNKDLLPNPPKTWEEIPALDKELKAKGKSALMFNLQEPYFTW
PLIAADGGYAFKYENGKYDIKDVGVDNAGAKAGLTFLVDLIKNKHMNADTDYSIAEAAFNKGETAMTINGPWAWSNIDT
SKVNYGVTVLPTFKGQPSKPFVGVLSAGINAASPNKELAKEFLENYLLTDEGLEAVNKDKPLGAVALKSYEEELAKDPR
IAATMENAQKGEIMPNIPQMSAFWYAVRTAVINAASGRQTVDEALKDAQTNSSSNNNNNNNNNNLGIEGRISTSGSGG
GGGSMSENLYFQGSMGVQVETISPGDGRTFPKRGQTCVVHYTGMLEDGKKFDSSRDRNKPFKFMLGKQEVIRGWE
EGVAQMSVGQRAKLTISPDYAYGATGHPGIIPPHATLVFDVELLKLNEGGSGGSGGSGGSMVSKGEELFTGVVPILVEL
DGDVNGHKFSVSGEGEGDATYGKLTLKFICTTGKLPVPWPTLVTTLTYGVQCFSRYPDHMKQHDFFKSAMPEGYVQE
RTIFFKDDGNYKTRAEVKFEGDTLVNRIELKGIDFKEDGNILGHKLEYNYNSHNVYIMADKQKNGIKVNFKIRHNIEDGS
VQLADHYQQNTPIGDGPVLLPDNHYLSTQSALSKDPNEKRDHMVLLEFVTAAGITLGMDELYKSGRHHHHHH

#### 4. Computational details

##### Myoglobin backbone idealization with inpainting

The backbone idealization of Rosetta-relaxed crystal structure of human myoglobin (PDB: 3RGK) was performed using RoseTTAFold joint inpainting.[10] Two separate design trajectories were performed. In first, the following regions were considered for idealization: 9 N-terminal residues, 10 C-terminal residues, positions 73-88 connecting the E and F helices. In the second strategy, in addition to the above, also positions 47-59 in the CD-loop region were considered for remodeling. Furthermore, positions in the fixed parts of the protein that are in contact with the remodeled regions and are not part of the heme binding site were allowed to be redesigned using the "inpaint_seq" option.

The following settings were included in the input JSON files to perform the design:

<u>Strategy 1:</u>

```
[{"pdb": "../3RGK_fr.pdb",
"task": "hal",
"dump_all": true,
"inf_method": "multi_shot",
"n_cycle": 15,
"num_designs": 20,
"tmpl_conf": "0.9",
"contigs": ["6-10,A10-72,14-19,A89-139,8-12"],
"inpaint_seq": ["A130","A134","A137"],
"out": "3RGK_inpaint1"}]
```

<u>Strategy 2:</u>

```
[{"pdb": "../3RGK_fr.pdb",
"task": "hal",
"dump_all": true,
"inf_method": "multi_shot",
"n_cycle": 15,
"num_designs": 10,
"tmpl_conf": "0.9",
"contigs": ["6-10,A10-46,10-16,A60-72,14-19,A89-139,8-12"],
"inpaint_seq": ["A26","A30","A34","A62","A130","A134","A137"],
"out": "3RGK_inpaint2"}]
```

##### ProteinMPNN design of myoglobin

The following command was used to perform ProteinMPNN[11] sequence redesign of the native myoglobin as well as the structures obtained from inpainting backbone idealization.

```
python $MPNN_PATH/protein_mpnn_run.py --jsonl_path ../parsed_pdbs_bb.jsonl
--fixed_positions_jsonl ../masked_pos.jsonl --batch_size 1 --out_folder ./
```

```
--num_seq_per_target 20 --sampling_temp "0.1 0.2 0.3" --omit_AAs='MC'
--checkpoint_path $MPNN_PATH/vanilla_model_weights/v_48_020.pt
```

Where `parsed_pdbs_bb.jsonl` contains the parsed PDB file information, created with the script `$MPNN_PATH/helper_scripts/parse_multiple_chains.py`

`masked_pos.jsonl` file contains the positions that were kept fixed during sequence design:

```
{"3RGK": {"A": [39, 42, 43, 45, 64, 67, 68, 71, 72, 89, 92, 93, 97,
99, 104, 107, 138]}}
```

For each of the outputs from the inpainting backbone idealization, the fixed position numbers were readjusted to correspond to the positions in the parent structure.

**ProteinMPNN design of TEV protease**

The following command was used to perform sequence design with ProteinMPNN on TEV protease.

```
python $MPNN_PATH/protein_mpnn_run.py \
        --jsonl_path ../parsed_pdbs_bb.jsonl \
        --chain_id_jsonl ../assigned_chains.jsonl \
        --fixed_positions_jsonl ../masked_pos.jsonl \
        --out_folder $MPNN_OUTDIR \
        --num_seq_per_target 16 \
        --sampling_temp "0.1 0.2 0.3" \
        --batch_size 8 \
        --omit_AAs='XC'
```

Where `../assigned_chains.jsonl` contains the parsed PDB chain information: `{"TEVd": [["A"]] }`

Sets of designs were distinguished by selection of fixed residues.

Designs with only the amino acid identities of the active site fixed during sequence design had the following residues fixed:

```
[31, 32, 44, 46, 81, 134, 135, 139, 146, 147, 148, 149, 150,151, 167, 168, 169,
170, 171, 172, 173, 174, 175, 176, 177, 178,204, 208, 209, 211, 213, 214, 215,
216, 217, 218, 219, 220]
```

Designs with the amino acid identities of active site residues and the 30% most conserved residues fixed during sequence design had the following residues fixed:

```
[3, 7, 9, 10, 11, 12, 14, 19, 25, 34, 36, 38, 42, 44, 46, 47, 48, 51, 52, 53,
55, 61, 62, 64, 68, 81, 88, 89, 90, 92, 94, 100, 101, 103, 110, 113, 116, 117,
126, 127, 129, 139, 140, 142, 143, 144, 146, 149, 151, 152, 154, 156, 160, 161,
163, 165, 167, 169, 177, 186, 190, 198, 202, 211, 212, 221]
```

Designs with the amino acid i2dentities of active site residues and the 50% most conserved residues fixed during sequence design had the following residues fixed:

```
[2, 3, 7, 8, 9, 10, 11, 12, 13, 14, 21, 23, 25, 26, 27, 31, 32, 34, 35, 36, 37,
38, 41, 42, 43, 44, 46, 47, 48, 51, 52, 53, 55, 59, 61, 62, 64, 68, 70, 72, 76,
81, 85, 88, 89, 90, 91, 92, 93, 94, 95, 98, 100, 101, 103, 107, 109, 112, 113,
115, 116, 117, 119, 123, 125, 126, 127, 129, 133, 134, 135, 139, 140, 141, 142,
143, 144, 146, 147, 148, 149, 150, 151, 152, 153, 154, 156, 157, 160, 161, 163,
165, 167, 168, 169, 170, 171, 172, 173, 174, 175, 176, 177, 178, 179, 182, 183,
186, 190, 198, 200, 202, 204, 205, 208, 209, 211, 212, 213, 214, 215, 216, 217,
218, 219, 220, 221]
```

Designs with the amino acid identities of active site residues and the 70% most conserved residues fixed during sequence design had the following residues fixed:

```
[1, 2, 3, 4, 7, 8, 9, 10, 11, 12, 13, 14, 15, 18, 21, 22, 23, 25, 26, 27, 31,
32, 33, 34, 35, 36, 37, 38, 40, 41, 42, 43, 44, 46, 47, 48, 49, 50, 51, 52, 53,
55, 57, 59, 61, 62, 63, 64, 66, 68, 69, 70, 71, 72, 73, 76, 79, 80, 81, 83, 84,
85, 86, 87, 88, 89, 90, 91, 92, 93, 94, 95, 96, 97, 98, 100, 101, 103, 107,
108, 109, 111, 112, 113, 115, 116, 117, 118, 119, 120, 122, 123, 124, 125, 126,
127, 129, 131, 133, 134, 135, 137, 139, 140, 141, 142, 143, 144, 145, 146, 147,
148, 149, 150, 151, 152, 153, 154, 155, 156, 157, 158, 160, 161, 163, 164, 165,
166, 167, 168, 169, 170, 171, 172, 173, 174, 175, 176, 177, 178, 179, 182, 183,
186, 187, 189, 190, 194, 196, 198, 200, 202, 203, 204, 205, 206, 207, 208, 209,
211, 212, 213, 214, 215, 216, 217, 218, 219, 220, 221]
```
